# Longitudinal evidence for the emergence of multiple intelligences in assistance dog puppies

**DOI:** 10.1101/2024.09.26.615218

**Authors:** Hannah Salomons, Morgan Ferrans, Candler Cusato, Kara Moore, Vanessa Woods, Emily Bray, Brenda Kennedy, Theadora Block, Laura Douglas, Ashton Roberts, Margaret Gruen, Brian Hare

## Abstract

Cognitive test batteries suggest that adult dogs have different types of cognitive abilities that vary independently. In the current study, we tested puppies repeatedly over a crucial period of development to explore the timing and rate at which these different cognitive skills develop. Service dog puppies (n = 113), raised using two different socialization strategies, were either tested longitudinally (n =91) or at a single time point (n = 22). Subjects tested longitudinally participated in the battery every two weeks during and just beyond their final period of rapid brain growth (from approximately 8-20 weeks of age). Control puppies only participated in the test battery once, which allowed us to evaluate the impact of repeated testing. In support of the multiple intelligences hypothesis (MIH), cognitive skills emerged at different points across the testing period, not simultaneously. Maturational patterns also varied between cognitive skills, with puppies showing adult-like performance on some tasks only weeks after a skill emerged, while never achieving adult performance in others. Differences in rearing strategy did not lead to differences in developmental patterns while, in some cases, repeated testing did. Overall, our findings provide strong support for the MIH by demonstrating differentiated development across the cognitive abilities tested.

---

Domestic dogs have a range of cognitive skills they use to flexibly solve problems in the physical and social world (Bräuer & Kaminski, 2021; Hare & Ferrans, 2021; Hare & Woods, 2013; Kaminski & Marshall-Pescini, 2014; Miklósi, 2014). Cognitive test batteries provide evidence that dogs have different types of cognitive abilities that vary independently. When large numbers of dogs (N>500) are tested across a range of different problems, individual skill for solving related cognitive tasks often correlate. However, performance on tasks designed to recruit different forms of cognition remains largely uncorrelated (Gnanadesikan et al., 2020; Horschler et al., 2019; MacLean, Herrmann, et al., 2017; Stewart et al., 2015; although see Bray et al, 2014). Performance within and across individuals is also stable on several cognitive measures when tested longitudinally as part of a larger test battery. Puppies who were above or below average at 10 weeks tend to maintain their position on a subset of tasks relative to other dogs when tested again as two-year-old adults (Bray, Gruen, et al., 2021).

These individual differences are relevant in the real world. Cognitive performance on a range of social and non-social tasks is associated with and predictive of training performance in working dogs (Bray et al., 2019; Bray, Otto, et al., 2021; Bray, Sammel, Cheney, et al., 2017; Hare & Ferrans, 2021; Lazarowski et al., 2019, 2020; MacLean & Hare, 2018). Cognitive differences across individuals have also been linked to training success and unwanted behaviors in pet dogs (Junttila et al., 2024). These same individual differences may be targeted with selection since they are heritable (with cognitive abilities previously identified as independent also varying in their degrees of heritability) (Bray, Gnanadesikan, et al., 2021; Gnanadesikan et al., 2020).

This body of work provides strong support for the hypothesis that adult dogs have multiple types of intelligence (Hare & Ferrans, 2021), which we will refer to as the multiple intelligences hypothesis (MIH). The MIH stems from the idea that evolution shapes organisms to solve the problems they are most likely to face in their environments in ways that promote their survival and reproduction (Hare, 2001; Hare & Wrangham, 2002; Tomasello, 1997). Different cognitive abilities provide different forms of flexible problem solving to overcome a range of novel challenges an organism predictably encounters (e.g. finding food, competing for mates, cooperating, etc.; (Shettleworth, 2009; Tomasello, 2022). The MIH does not preclude a hierarchical structure in which variation in different cognitive skills is, in part, explained by a higher order cognitive variable(s) (i.e. Bognár et al., 2024; Herrmann et al., 2010; Mithen, 1996; Sternberg, 1999). However, it does stand in stark contrast to any hypothesis positing that a single, general form of intelligence or unidimensional cognitive factor such as “learning ability” can explain cognitive diversity across individuals and species (Bastos & Taylor, 2020; Deaner et al., 2007; Macphail & Bolhuis, 2001; Shettleworth, 2009; Tomasello et al., 1997). We will call this the general intelligence hypothesis (GIH).

Cognitive ontogeny provides a critical test of these two models of cognitive architecture. The MIH predicts cognitive abilities should show differentiated developmental patterns (Herrmann et al., 2010; Wobber et al., 2014). As the brain develops, the different forms of cognition it supports should emerge separately. The timing of emergence and maturation of cognitive abilities are predicted to differ, in part, because brain areas that support different types of cognition can develop at different rates. Alternatively, the GIH predicts that cognitive abilities should show a unified developmental pattern. Emergence and development could vary across individuals, based largely on experiential differences, but no clear order of cognitive emergence should be observed across a group between different cognitive tasks, and no maturational pace should be specific to a certain cognitive task. Therefore, finding differentiated development patterns in would provide a powerful falsification of the GIH.

Developmental comparisons provide preliminary support for the MIH. Compared to wolf puppies, dog puppies show early emerging use of human gestures and eye contact but similar memory and self-control abilities (Salomons et al., 2021). Lazarowski et al (2020) tested a group of dogs raised, bred and trained for detection work using a cross-sectional design, and found that dogs were skilled by six months in a self-control task while skill with a delayed memory task developed in the eleven-month-old puppies. While these previous studies support the developmental prediction of the MIH, they have relatively low resolution. They only test puppies at a single time point during their early development.

By eight weeks a puppy is showing some key features of independence, such as being weaned and walking, but its brain is still rapidly developing until at least 16 weeks of age(Gross et al., 2010). Dog brains, like those of humans, rely on white matter to rapidly communicate signals between areas of the brain. The relative white to grey matter intensity and myelination of critical neuron networks begins approaching adult levels by 16 weeks. The full length of the corpus collosum, or the part of the brain that connects the two hemispheres of the brain and allows them to communicate, also reaches adult form around 16 weeks of age (Gross et al., 2010; Schmidt et al., 2012). While a puppy’s brain continues to develop after 16 weeks, the period between weaning and 16 weeks represents a final period of rapid brain growth (i.e. the first postnatal stage of rapid growth occurs during nursing). It is during this period that different cognitive abilities are most likely to first emerge, but detecting their emergence will require a higher frequency of testing than previous studies (Bray et al., 2020; Bray, Gruen, et al., 2021; Lazarowski et al., 2020; Salomons et al., 2021).

To more fully test the MIH and document cognitive development, what is needed is a high-resolution longitudinal approach that repeatedly samples puppies during their final period of rapid brain growth between 8-16 weeks of age. Here we provide a critical test of this core developmental prediction of the MIH. We measure a range of cognitive abilities in assistance dog puppies previously shown to be critical to their training success as adults. To test the impact of socialization during rearing on cognitive maturation rates we compared two groups of puppies raised in different ways. To examine the role of repeated testing on performance we compared the final test session of our longitudinal sample to an age matched control group only tested once. The MIH predicts that the different cognitive abilities will emerge and mature at different times and paces during this period of rapid brain growth. While rearing and repeated testing may impact development, a differentiated maturational pattern will still be discernable. In contrast, the GIH suggests that improvement across tasks will be uniform and tightly linked to rearing experience or exposure to each task. After accounting to for rearing and test exposure, individual cognitive skills will not show significant group level patterns of emergence and maturation related to their age.

## Methods

### Subjects

113 (M:F = 57:56) individual puppies participated in the Dog Cognitive Longitudinal Battery (DCLB) and are included in this analysis. Longitudinal subjects (*n = 91*) were tested on the battery multiple times between 8-20 weeks of age. Control subjects (*n = 22*) completed the battery once, typically at 16-20 weeks of age (2 controls were tested at 12-13 weeks of age due to the pandemic closures, Table 1). All puppies were Labrador retrievers, golden retrievers, or Labrador-golden retriever crosses with known pedigrees from lines of dogs being bred and socialized for work as assistance dogs (Bray, Gruen, et al., 2021). The majority of puppies came from Canine Companions (CC, www.canine.org, *n = 97*), while the remainder came from Eyes, Ears, Nose, and Paws (EENP, www.eenp.org, *n =* 14), and Guiding Eyes for the Blind (GEB, www.guidingeyes.org, *n =* 2).

**Table 1:**
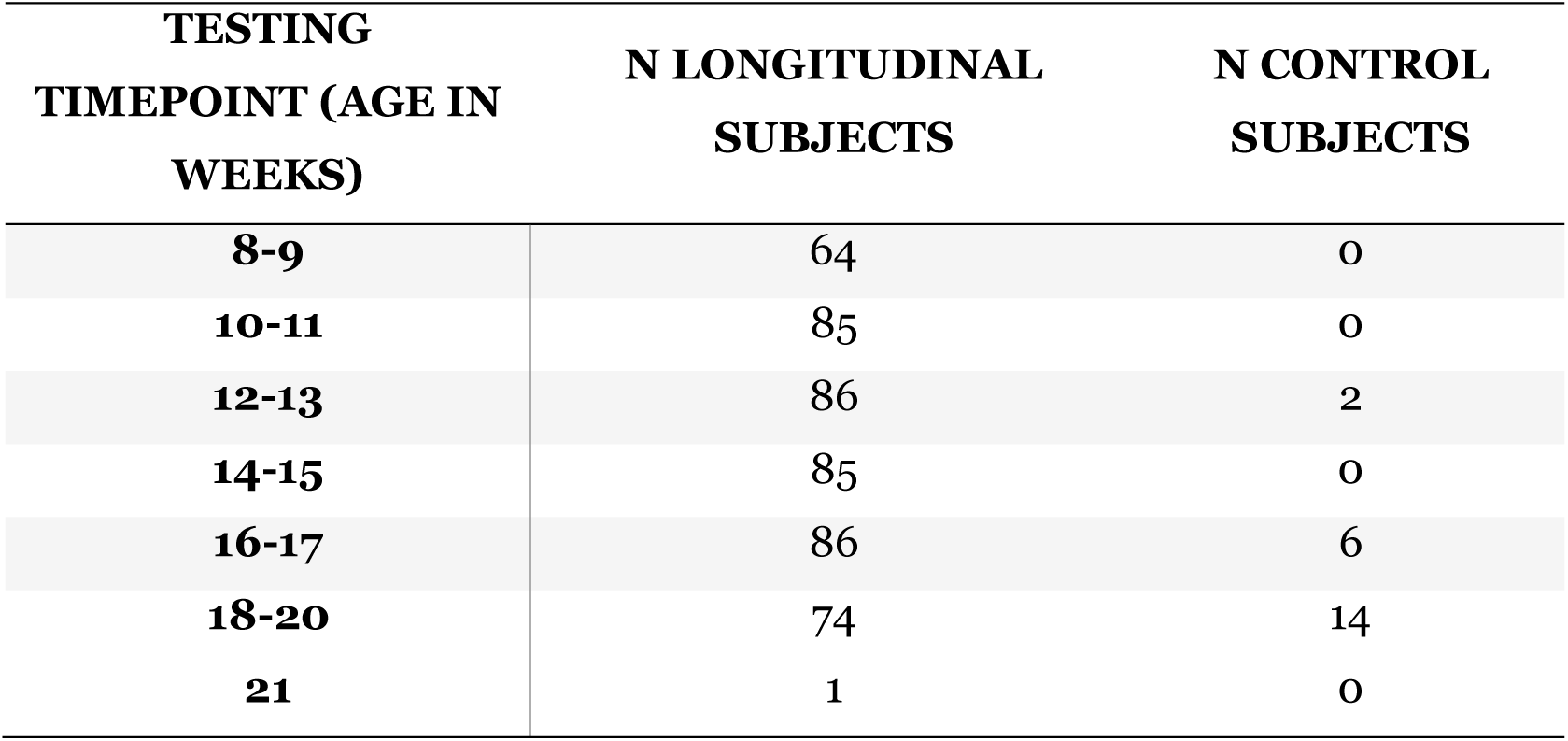
Number of subjects tested on the DCLB at each timepoint. 91 individuals were tested longitudinally (i.e., across multiple timepoints), and 22 individuals were tested at a single timepoint as controls.

Puppies were whelped and reared with their mothers and littermates until weaning at 6-8 weeks of age, either at a whelping center or in a volunteer’s home, according to each organization’s protocols. After weaning, and by the time of testing, all puppies were reared in one of two conditions: in a raiser’s family home (intensive socialization strategy, *n =* 78), or in the Duke Puppy Kindergarten (DPK) (extensive socialization strategy, *n =* 35). In both rearing conditions, puppies were raised according to the guidelines of the organization to which they belonged. Within these guidelines their socialization to humans and same-aged puppies differed. Intensively socialized home-raised puppies primarily interacted with their raiser families, and were the only puppy in the home. Typically, their raisers introduced them to new places and people on a weekly basis, and socialized them with other similarly aged puppies at monthly training classes. The extensively socialized DPK-raised puppies were cared for by dozens of individuals each day. This included 1-7 caretakers at a time during the day, who rotated every 2-4 hours from a team of over 100 undergraduate student volunteers. DPK puppies also spent significant time daily with the other same-aged puppies in the program. At night, DPK puppies were cared for by a smaller cohort of student volunteers, either in their homes/dorms or DPK’s husbandry room, who rotated every 2-3 nights. Additionally, they went on daily outings around Duke’s campus and participated in public visiting hours most days, resulting in interactions with hundreds of new people in total during their time at DPK. Our extensive and intensive rearing approaches were designed after contacting six representative assistance dog organizations and reviewing their puppy raising manuals. While both consist of highly positive socialization, the intensive raising strategy was more typical of a pet dog’s experience, while the extensive strategy was more of an extreme in terms of the exposure to humans and other puppies among the different strategies we surveyed. While representing very different approaches to socialization, each may have advantages in increasing bonding, positive social behaviors, and learning outcomes (Harvey et al., 2016; Vaterlaws-Whiteside & Hartmann, 2017).

Rearing group assignments were based on the logistics of volunteer availability and timing of whelping. When possible, preference for participation in the study was given to non-littermates, with the aim of including puppies from as many unique litters as possible. The 113 puppies were conceived from 69 unique dam/sire pairings, born between November 2018 and November 2023. Approximately 65% (*n =* 74) of subjects had a full sibling, and ∼31% (*n =* 35) had a half sibling in the sample. Useable data were obtained from all subjects, although not all subjects completed all trials of every test session. Incomplete testing occurred occasionally due to puppies’ inability to finish testing (see abort criteria) or logistical constraints (e.g. pandemic closures, illness, age-constrained timing not coinciding with staff availability, etc.). See **Table 1** for the total number of subjects in the longitudinal and control sample at each age.

### Procedure

#### Setup of Testing Area

All experiments took place from October, 2019 to November, 2023 in a quiet testing room at either the Duke Canine Cognition Center, or one of Canine Companions’ campuses. To further reduce any distractions, a white noise machine was used during all testing. The testing area was approximately 8’ x 10’ with a 4’ x 6’ testing mat centered inside (**Figure 1**). A short fence was placed around the testing area to help limit distractions and allow puppies to focus on the experimental manipulations and rewards. Water was available *ad libitum* via a bowl of fresh water placed in the testing space behind the start box (see **Figure 1**). All testing sessions were filmed with cameras positioned to capture the behaviors of the subject and experimenter for each test. Depending on the test, this required two or three video cameras (Sony Handycam HDR-CX405 or similar). Cameras were always mounted on tripods and positioned outside the testing area.

**Figure 1:**
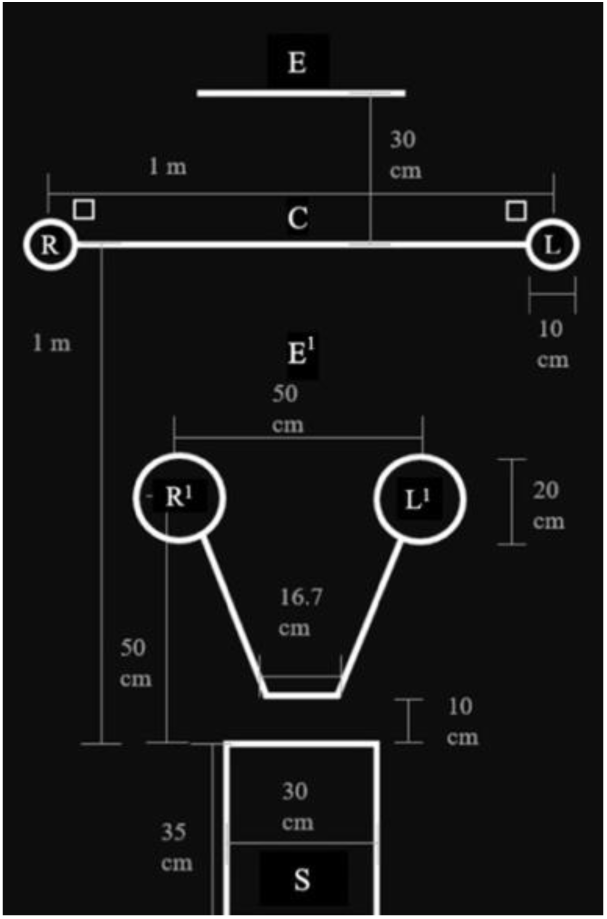
DCLB Cognitive Testing Mat. Letters indicate key locations: puppy start box (S), experimenter positions (E, E^1^), and possible bait locations (R, C, L, R^1^, L^1^).

### General Methods

Subjects were given a few minutes to explore and acclimate to the testing room and experimenters before testing began. An experimenter (E) conducted the tests and a handler (H) gently positioned and held the subject in the starting position (i.e. in the starting box; see Figure 1). Subjects began all test trials sitting in the starting box facing E. H sat or kneeled behind the subject, gently holding them in position either by the collar or by placing a hand on either side of the chest. E would gain the subject’s attention, carry out the experimental manipulation, and then indicate to H to release the subject by saying “Okay”. In timed trials, H would also start the timer once E said “Okay”. H released the subject by gently removing their hand(s), making sure to remove both hands evenly to avoid biasing the subject’s direction. To limit unintentional influence on subject’s choices, when not conducting the experimental manipulation, E assumed a “resting position” where she kneeled and looked down at her lap/knees with her hands behind her back. Whenever possible, the same person played the role of E for a given puppy across sessions. However, different people played the roles of E and H across puppies.

In each test, subjects were trying to obtain food rewards. One piece of puppy kibble was typically used for each trial’s reward. Subjects were given a set amount of time to search for the reward or otherwise participate in each trial (see procedures described for each test below). When a subject did not participate before the time expired (“no-choice”) twice in a row, they were refamiliarized with the test procedure. If the puppy did not engage in refamiliarization trials, made a total of four no-choices, or at any time exhibited signs of significant stress (including excessive whining, barking, escape behavior, or defecation), the session was aborted and attempted again after a break of at least 30 minutes. Additional task-specific choice and abort criteria are described in the supplemental methods section.

Cognitive tasks in the DCLB were chosen based on those of previous research batteries (DCTB, MacLean, Herrmann, et al., 2017; DCDB, Bray et al., 2020a) which were associated with training success and/or exhibited significant development or individual stability from puppyhood to adulthood (Bray et al., 2019; Bray, Gruen, et al., 2021; Bray, Otto, et al., 2021; MacLean & Hare, 2018;**Table S1**). More information comparing the DCLB with the DCDB is contained in the supplemental methods.

The DCLB consists of 2 warmup sessions, 9 cognitive tasks, and 4 temperament measures as part of a larger longitudinal research program. To test the predictions of our hypotheses, only the cognitive tasks are reported on in this study. Cognitive testing took place over two sessions, which were completed across one or two days based on puppy and raiser availability, with temperament tasks typically being run between the two sessions.

**Table 2:** Order of DCLB Cognitive Testing Tasks.

| **Session 1** | **Session 2** |
| --- | --- |
| Unsolvable (Eye Contact) | Cylinder (Inhibitory Control & Reversal Learning) |
| Bowl Choice Warm-Ups | Bowl Choice Warm-Ups |
| Marker Gesture | Momentary Pointing Gesture |
| Working Memory (Distraction) | Working Memory (Delay) |
| Causal Reasoning | Auditory Discrimination |
|  | Odor Discrimination |

### Design

This study has a within-subject, longitudinal design. The study was run as a battery of tests, in which subjects participated in all the cognitive tests in the same order. Longitudinal subjects were tested on the DCLB every two weeks for ten weeks, from ages 8-18, 9-19, or 10-20 weeks (typically completing 6 test sessions, though some participants missed sessions due to availability constraints). To assess the impact of repeated testing on the longitudinal sample, control subjects were tested on the DCLB only once. In tasks where food was hidden in one of two locations, the order across trials of the “correct” baited location was the same for all subjects (following Herrmann et al., 2007; MacLean & Hare, 2018). Reward placement was pseudo-randomized across the different tasks in the battery using the constraint that the reward was hidden on the right and left an equal number of times in each task and never on the same side more than twice in a row. The dependent variable for each test was generally whether or not the puppy correctly solved the test (with the exception of Unsolvable which measured seconds of eye contact), and are described in detail in the supplemental methods section. The independent variables of interest include age (in weeks), rearing strategy (DPK vs home-raised), and prior testing experience (number of prior testing sessions).

### Scoring and Analysis

Choice and scoring definitions for each test are included in the supplemental methods section. All trials were videotaped from an angle (or multiple angles) which captured the subject’s response as well as the experimenter (E). Subject responses were live-coded by E after each trial for all cognitive measures. In a minority of cases E was unable to live-code (e.g. due to stopwatch malfunction) and later coded the trial from video. Approximately 20% of all sessions (chosen pseudo-randomly ensuring that subjects from both rearing strategies as well as controls were included) were independently coded for reliability from video. Reliability was excellent on all tests: Marker Gesture (Cohen’s κ = 0.98, *n =* 405, p <.001); Momentary Pointing (Cohen’s κ = .95, *n =* 396, p <.001); Working Memory (Delay) (Cohen’s κ = .99, *n =* 410, p <.001); Working Memory (Distraction) (Cohen’s κ = .98, *n =* 400, p <.001); Causal Reasoning (Cohen’s κ =.97, *n =* 397, p <.001); Auditory Discrimination (Cohen’s κ = .97, *n =*397, p <.001); Odor Discrimination (Cohen’s κ = .96, *n =* 390, p <.001); Cylinder (Inhibitory Control) (Cohen’s κ = .96, *n =388*, p <.001); Cylinder (Reversal Learning) (Cohen’s κ = .94, *n =386*, p <.001); Unsolvable (eye-contact) (Pearson’s *r*(392) = .96, p<.001).

Analysis was completed in RStudio v. 2023.06.1+524. Mixed effects logistic and linear regression models (depending on the test -logistic for binary dependent variables and linear for continuous) were run, including age, rounds of previous test battery experience, and rearing strategy (DPK vs home-raised) as variables. These models were compared using AIC scores to determine which variables significantly impacted the outcome of each test and the nature of their effects. Age and experience can be differentiated in these analyses, in part, because the start of biweekly testing was not perfectly synchronized across subjects. Some puppies started testing when they were either eight (n=32), nine (n=30) or ten-fourteen weeks (n=29) of age and were tested biweekly from their first test date (**see Table S14**). In addition, some puppies did not complete all testing within each session (e.g. following the abort criteria) or due to constraints on availability were tested on fewer than 6 longitudinal time points. Together, this means puppies of the same age did not always have the same amount of experience. This provided the opportunity to examine the influence of age and experience separately even within the longitudinal sample.

As another way to examine the effect of repeated testing on performance, two-tailed Welch’s t-tests were conducted to compare the performance of longitudinal subjects in their final (fifth or sixth) testing session with same-age (17-21 weeks) control subjects in their sole testing session (longitudinal mean age = 18.52 weeks; control mean age = 18.55 weeks; two-tail Welch’s t-test shows no significant difference in age, t(25.07) = 0.10, p=.92). To determine age of emergence of each skill, Z tests were run for each age group against the null hypothesis of 50% correct (intercept = 0) using a mixed effect logistic regression model.

For the skills on which puppies reached above chance performance at any point during the 8-21 week age window and which also showed a positive linear relationship with age, segmented regression analysis was performed and a breakpoint was determined using the Segmented package (Muggeo, 2008) with age as the independent variable and proportion of trials correct was the dependent variable. The breakpoint is defined as the age at which there is a significant change in the slope of the performance in relation to age.

### DCLB Task Protocols

In the marker gesture task, the experimenter hides food in one of two bowls (placed in positions R and L, **Figure 1**) behind an occluder. Once the occluder is removed and the bowls are revealed, the experimenter uses ostensive cues (i.e., attempts to make eye contact and says “puppy look!” in a high-pitched voice) while placing a small yellow wooden block next to the correct bowl to indicate the hiding location.

In the working memory tasks, the puppy is allowed to see which of the two bowls the food is placed into, and then released to search for it after 20 seconds. In the “delay” condition, the experimenter is still and silent during the delay; in the “distraction” condition, the experimenter engages the puppy in play with a toy during the delay.

In the auditory and odor discrimination tasks, the puppy is asked to determine which of the two close hiding locations (positions R^1^ and L^1^) contains the food based on these senses. In the auditory, a piece of dry kibble is audibly dropped into one of two metal bowls, while the other is silently sham-baited. In the odor task, two plastic tubes with openings covered by mesh fabric are presented to the puppy to sniff, only one of which contains food.

In the cylinder task, puppies retrieve food from a transparent cylinder – to succeed in the inhibitory control condition, they have to approach either open side of the cylinder rather than touch the cylinder walls through which they could see the food. In the reversal condition, the puppy’s preferred side from the previous trials is now blocked, and we observed if they could learn to approach the other side without bumping the cylinder.

In the unsolvable (eye contact) task, food is placed in a sealed transparent container, and the experimenter times how much eye contact the puppy makes with them while the food is inaccessible.

In the causal reasoning task, the experimenter places a cloth over a bowl containing food in one hiding location, while an identical cloth is laid flat on the floor in the other hiding location, and the puppy must choose a location.

The momentary pointing task is identical to the marker gesture task, except that instead of placing a marker, the experimenter indicates the hiding location by pointing with the contralateral arm, holds the point for three seconds, and then returns to resting position before the puppy is released to make a choice.

Detailed protocols for each task are provided in the supplementary materials.

## Results

### Marker Gesture Task

Age (in weeks), rearing strategy (DPK vs home-raised), and experience (number of prior test sessions completed) were considered as covariates for generalized linear mixed effects models (**Table S2**). Age was a significant predictor variable in all models it was considered in, while rearing strategy was never significant. Experience was a significant predictor variable when entered into the model alone, but became insignificant when added to models that also included age. The model with the lowest AIC score (876.116) contained only age as a predictor variable (β_age_ = 0.114, SE = 0.023, p<.001).

A two-tailed Welch’s t-test comparing the proportion of trials in which the baited bowl was chosen by the longitudinal subjects in their last testing session (only those aged 17-21 weeks old, *n =* 74, M ± SD = 0.91 ± 0.28) to that of control subjects in the same age range (*n =* 20, M ± SD = 0.91 ± 0.28) shows that they performed similarly (t(124.75) = 0.009, p=.99) (**Table 3**, **Figure S1**).

**Table 3:** Differences between longitudinal puppies in their final testing session and same age (17-21 weeks) control puppies. Bolded means for each measure indicate when a group performed significantly higher (note: eye contact measured in two ways in same task).

| Variable | Units | Longitudinal<br>Mean ± SD | Control<br>Mean ± SD | *t* | *df* | *p* |
| --- | --- | --- | --- | --- | --- | --- |
| Marker Gesture | Proportion<br>trials correct | 0.91 ± 0.28 | 0.91 ± 0.28 | 0.009 | 124.75 | .99 |
| Working Memory<br>(Delay) | Proportion<br>trials correct | 0.86 ± 0.35 | **0.95 ± 0.22** | 2.7208 | 176.78 | <.01 |
| Working Memory<br>(Distraction) | Proportion<br>trials correct | 0.79 ± 0.41 | 0.79 ± 0.41 | -0.06 | 124.08 | .95 |
| Auditory<br>Discrimination | Proportion<br>trials correct | 0.57 ± 0.50 | **0.69 ± 0.47** | 2.03 | 130.77 | <.05 |
| Odor<br>Discrimination | Proportion<br>trials correct | 0.50 ± 0.50 | 0.56 ± 0.50 | 0.90 | 123.05 | .37 |
| Cylinder<br>(Inhibitory<br>Control) | Proportion<br>trials correct | **0.94 ± 0.23** | 0.68 ± 0.47 | -4.87 | 89.99 | <.001 |
| Cylinder (Reversal<br>Learning) | Proportion<br>trials correct | **0.62 ± 0.49** | 0.28 ± 0.45 | -5.98 | 133.18 | <.001 |
| Unsolvable<br>(Looking Time) | No. of<br>seconds | **5.34 ± 6.50** | 3.07 ± 4.46 | -3.59 | 184.91 | <.001 |
| Unsolvable<br>(Prop. Looked) | Proportion<br>trials looked | **0.81 ± 0.39** | 0.60 ± 0.49 | -3.47 | 109.44 | <.001 |
| Causal Reasoning | Proportion<br>trials correct | **0.89 ± 0.31** | 0.43 ± 0.50 | -6.25 | 59.68 | <.001 |
| Momentary<br>Pointing | Proportion<br>trials correct | 0.51 ± 0.50 | 0.48 ± 0.50 | -0.45 | 117.66 | .65 |

Figure 2 (by weeks) and **Figure S2** (by days) graph all longitudinal puppies’ (*n =* 91) performance on the marker gesture task across the 8-21 week age window using the “geom_smooth” function, method = ‘loess’ (Wickham, 2016), and **Figure S3** shows the same grouped by rearing strategy.

**Figure 2:**
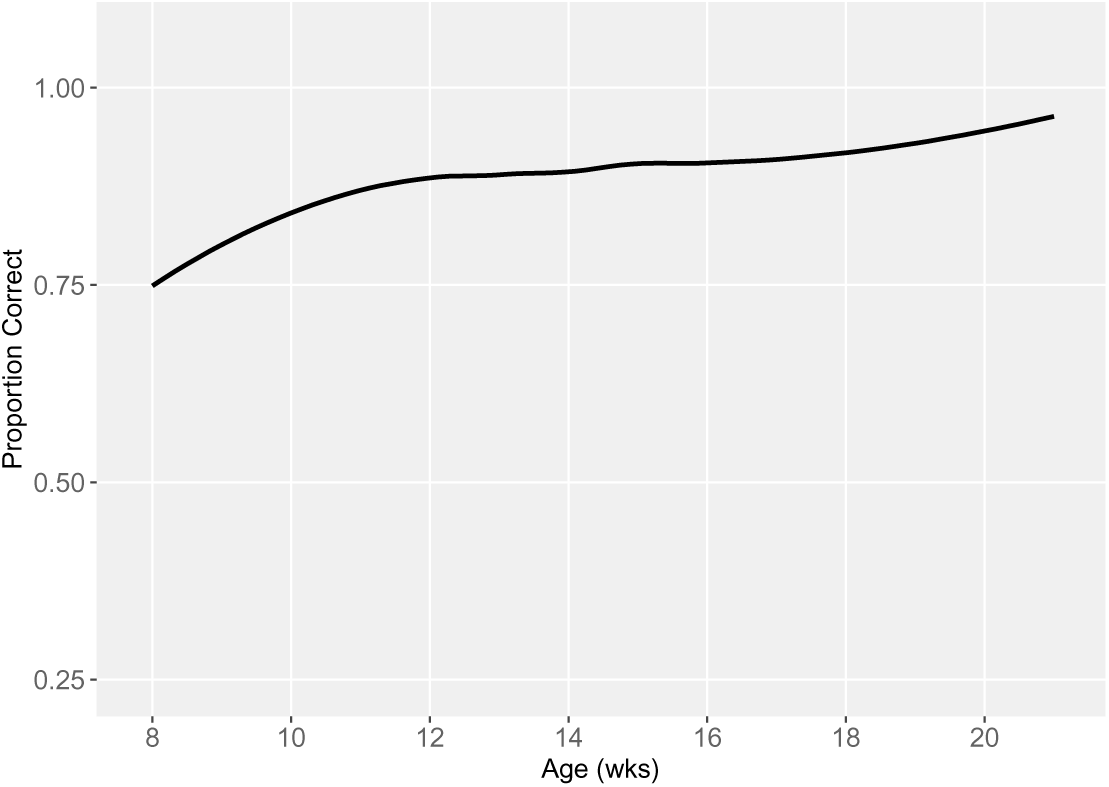
Performance on the Marker Gesture Task by Age in Weeks

### Working Memory (Delay) Task

Age (in weeks), rearing strategy (DPK vs home-raised), and experience (number of prior test sessions completed) were considered as covariates for generalized linear mixed effects models (**Table S3**). Age and Experience were significant predictor variables in all the models in which they were entered. Rearing was a significant predictor variable in the model in which it was entered alone and in the Age + Rearing model. The model with the lowest AIC score (1072.844) contained age and experience as predictor variables (β_age_ = 0.201, SE = 0.051, p<.001; β_experience_ =-0.300, SE = 0.103, p <.001).

A two-tailed Welch’s t-test comparing the proportion of trials in which the baited bowl was chosen by the longitudinal subjects on their last testing session (only those aged 17-21 weeks old, *n =* 76, M ± SD = 0.86 ± 0.35) to that of control subjects in the same age range (*n =* 19, M ± SD = 0.95 ± 0.22) shows that the controls performed significantly better than the longitudinal sample (t(176.78) = 2.7208, p<.01) (**Table 3**, **Figure S4**).

Figure 3 and **Figure S5** graph all longitudinal puppies’ (*n =* 91) performance on the working memory with delay task across the 8-21 week age window using the “geom_smooth” function, method = ‘loess’ (Wickham, 2016), and **Figure S6** shows the same grouped by rearing strategy.

**Figure 3:**
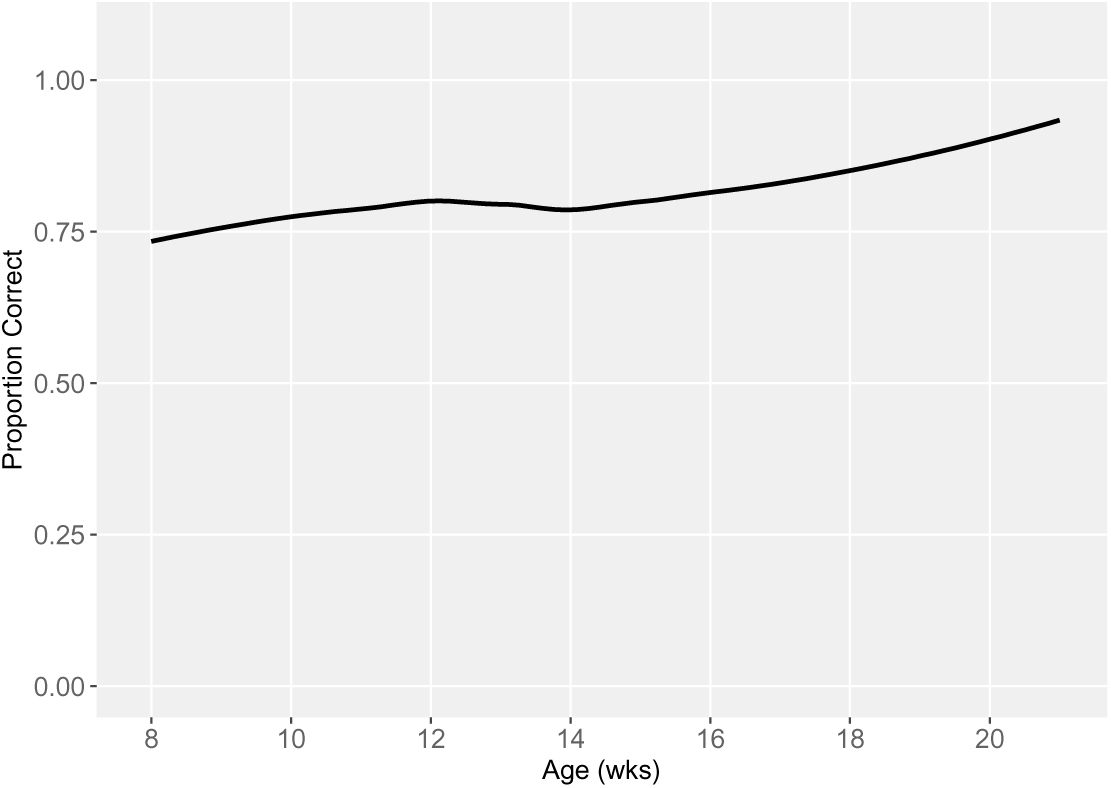
Performance on the Working Memory (Delay) Task by Age in Weeks

### Working Memory (Distraction) Task

Age (in weeks), rearing strategy (DPK vs home-raised), and experience (number of prior test sessions completed) were considered as covariates for generalized linear mixed effects models (**Table S4**). Age was a significant predictor variable in all models it was entered in. Experience was a significant predictor variable when entered into the model alone and with age, but not in the model with age and rearing. Rearing was not a significant predictor variable in any of the models in which it was entered. The model with the lowest AIC score (1222.845) was that which contained age and rearing as predictor variables (β_age_ = 0.091, SE = 0.016, p<.001; β_rearing_ =-0.311, SE = 0.204, ns).

A two-tailed Welch’s t-test comparing the proportion of trials in which the baited bowl was chosen by the longitudinal subjects on their last testing session (only those aged 17-21 weeks old, *n =* 74, M ± SD = 0.79 ± 0.41) to that of control subjects in the same age range (*n =* 20, M ± SD = 0.79 ± 0.41) shows that they performed similarly (t(124.08) = -0.06, p=.95) (**Table 3**, **Figure S7**).

Figure 4 and **Figure S8** graph all longitudinal puppies’ (*n =* 91) performance on the working memory with distraction task across the 8-21 week age window using the “geom_smooth” function, method = ‘loess’ (Wickham, 2016), and **Figure S9** shows the same grouped by rearing strategy.

**Figure 4:**
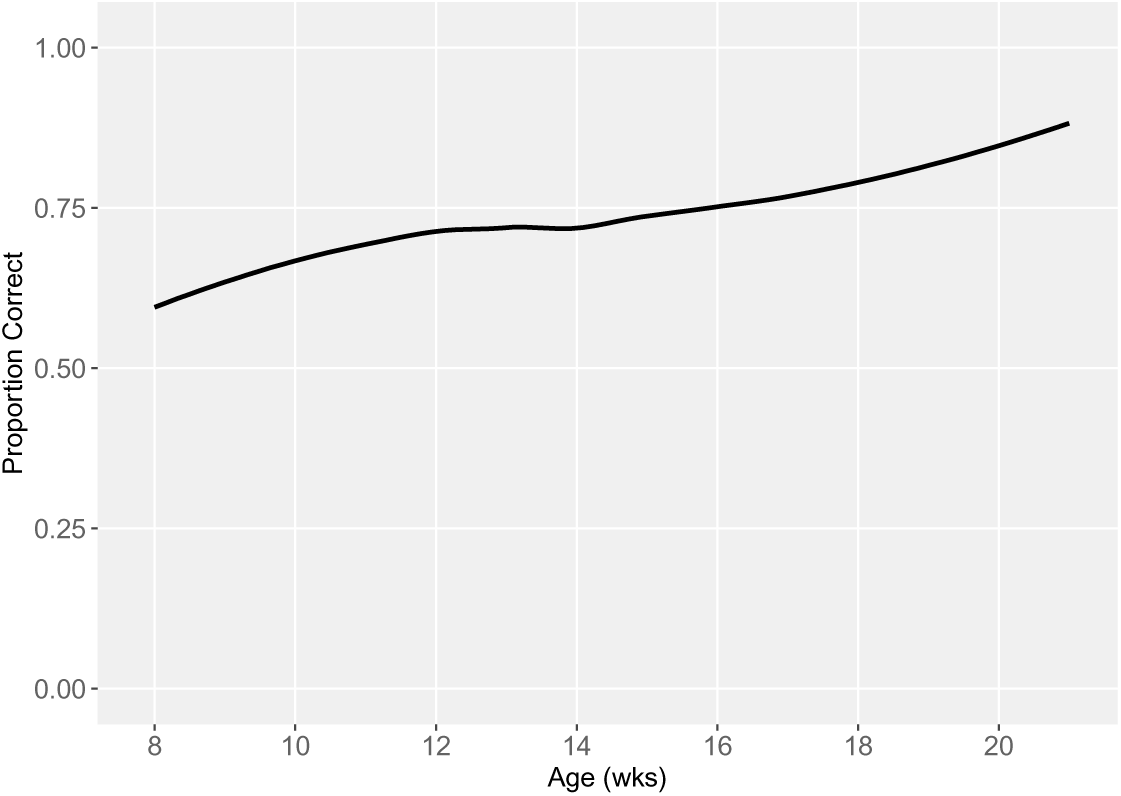
Performance on the Working Memory (Distraction) by Age in Weeks

### Auditory Discrimination Task

Age (in weeks), rearing strategy (DPK vs home-raised), and experience (number of prior test sessions completed) were considered as covariates for generalized linear mixed effects models (**Table S5**). Age was a significant predictor variable in the age alone model and the age + rearing model, but became insignificant when experience was entered into the models. Experience was a significant predictor in all the models in which it was entered. Rearing was not a significant predictor variable in any of the models in which it was entered. The model with the lowest AIC score (1205.470) contained only experience as a predictor variable (β_experience_= -0.135, SE = 0.029, p <.001).

A two-tailed Welch’s t-test comparing the proportion of trials in which the baited bowl was chosen by the longitudinal subjects on their last testing session (only those aged 17-21 weeks old, *n =* 75, M ± SD = 0.57 ± 0.50) to that of control subjects in the same age range (*n =* 20, M ± SD = 0.69 ± 0.47) shows that the control puppies performed significantly better than the longitudinal puppies (t(130.77) = 2.03, p<.05) (**Table 3**, **Figure S10**).

Figure 5 and **Figure S11** graph all longitudinal puppies’ (*n =* 91) performance on the auditory discrimination task across study period by weeks and days using the “geom_smooth” function, method = ‘loess’ (Wickham, 2016), and **Figure S12** shows the same grouped by rearing strategy.

**Figure 5:**
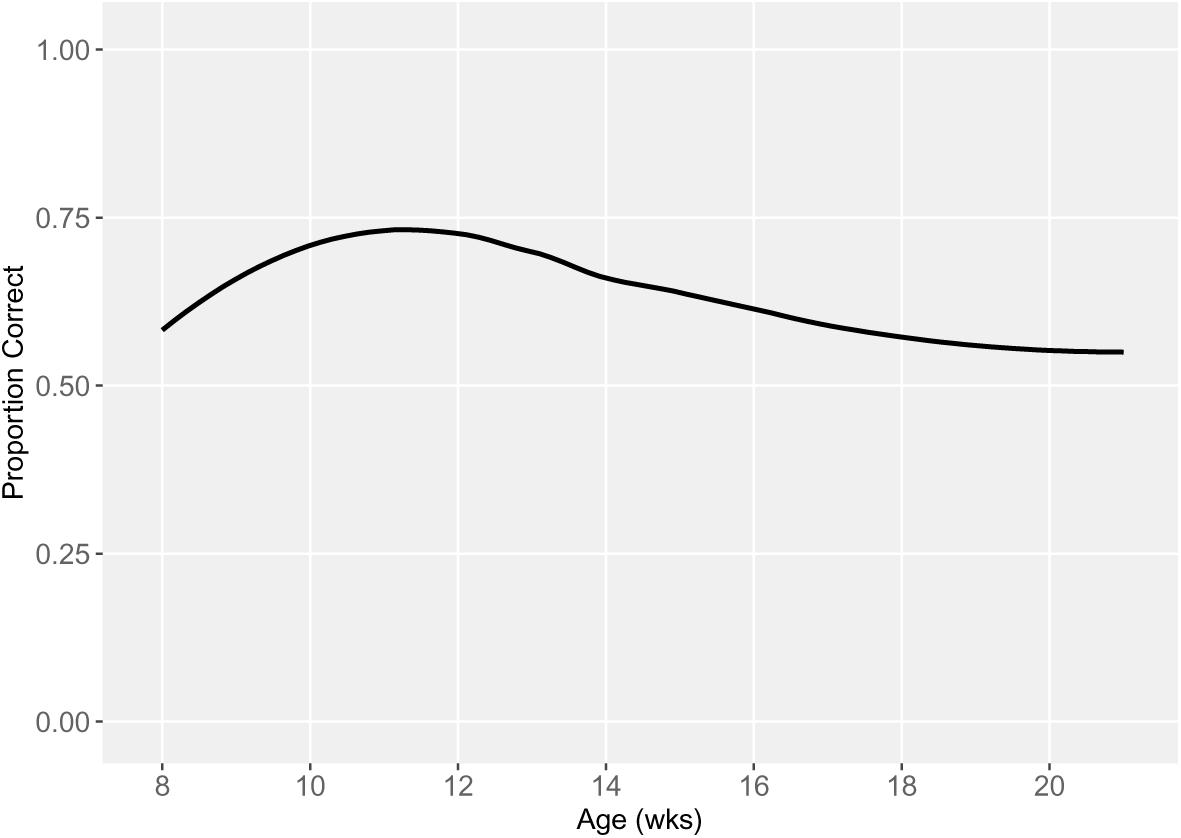
Performance on the Auditory Discrimination Task by Age in Weeks

### Odor Discrimination Task

Age (in weeks), rearing strategy (DPK vs home-raised), and experience (number of prior test sessions completed) were considered as covariates for generalized linear mixed effects models (**Table S6**). Age was a significant predictor variable in the age alone model and the age + rearing model. Experience was a significant predictor variable when entered into the model alone, but became insignificant when added to models that also included age. Rearing was not a significant predictor variable in any of the models in which it was entered. The model with the lowest AIC score (1,225.646) contained age and rearing as predictor variables (β_age_= -0.029, SE = 0.013, p<.01; β_rearing_=0.146, SE=0.099, ns).

A two-tailed Welch’s t-test comparing the proportion of trials in which the baited bowl was chosen by the longitudinal subjects on their last testing session (only those aged 17-21 weeks old, *n =* 74, M ± SD = 0.50 ± 0.50) to that of control subjects in the same age range (*n =* 19, M ± SD = 0.56 ± 0.50) shows that they performed similarly (t(123.05) = 0.90, p=.37) (Table 3, **Figure S13**).

Figure 6 and **Figure S14** graph all longitudinal (*n =* 91) puppies’ performance on the odor discrimination task across the 8-21 week age window using the “geom_smooth” function, method = ‘loess’ (Wickham, 2016), and **Figure S15** shows the same grouped by rearing strategy.

**Figure 6:**
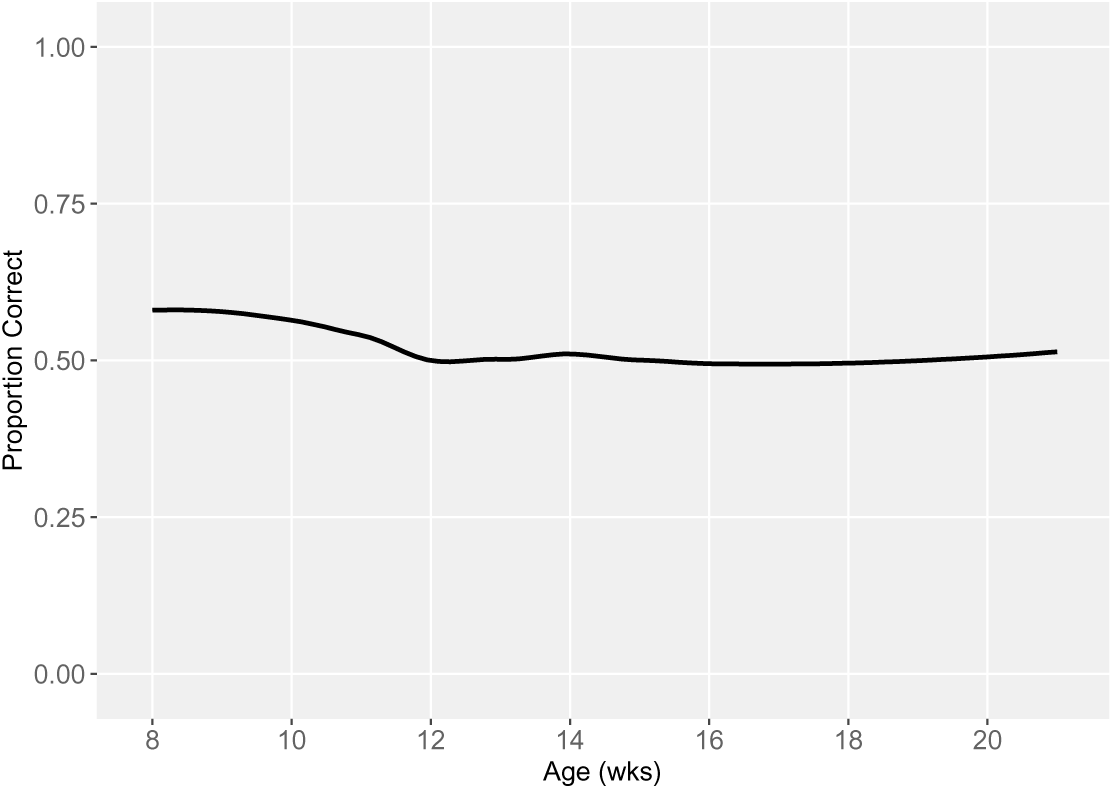
Performance on the Odor Discrimination Task by Age in Weeks

### Cylinder Task

#### Inhibitory Control trials

Age (in weeks), rearing strategy (DPK vs home-raised), and experience (number of prior test sessions completed) were considered as covariates for generalized linear mixed effects models (**Table S7**). Age and experience were both significant predictor variables in all the models in which they were entered. Rearing was not a significant predictor variable in any of the models. The model with the lowest AIC score (957.156) contained age and experience (β_age_= 0.084, SE = 0.028, p<.001; β_experience_ = 0.462, SE = 0.038, p<.001).

A two-tailed Welch’s t-test comparing the proportion of trials in which a clear route was taken to an open side without touching the transparent cylinder wall by the longitudinal subjects on their last testing session (only those aged 17-21 weeks old, *n =* 75, M ± SD = 0.94 ± 0.23) to that of control subjects in the same age range (*n =* 20, M ± SD = 0.68 ± 0.47) shows that the longitudinal puppies performed significantly better than the control puppies (t(89.99) = -4.87, p<.001) (Table 3, **Figure S16**).

Figure 7 and Figure S17 graph all longitudinal puppies’ (*n =* 91) performance on the inhibitory control trials across the 8-21 week age window using the “geom_smooth” function, method = ‘loess’ (Wickham, 2016), and **Figure S18** shows the same grouped by rearing strategy.

**Figure 7:**
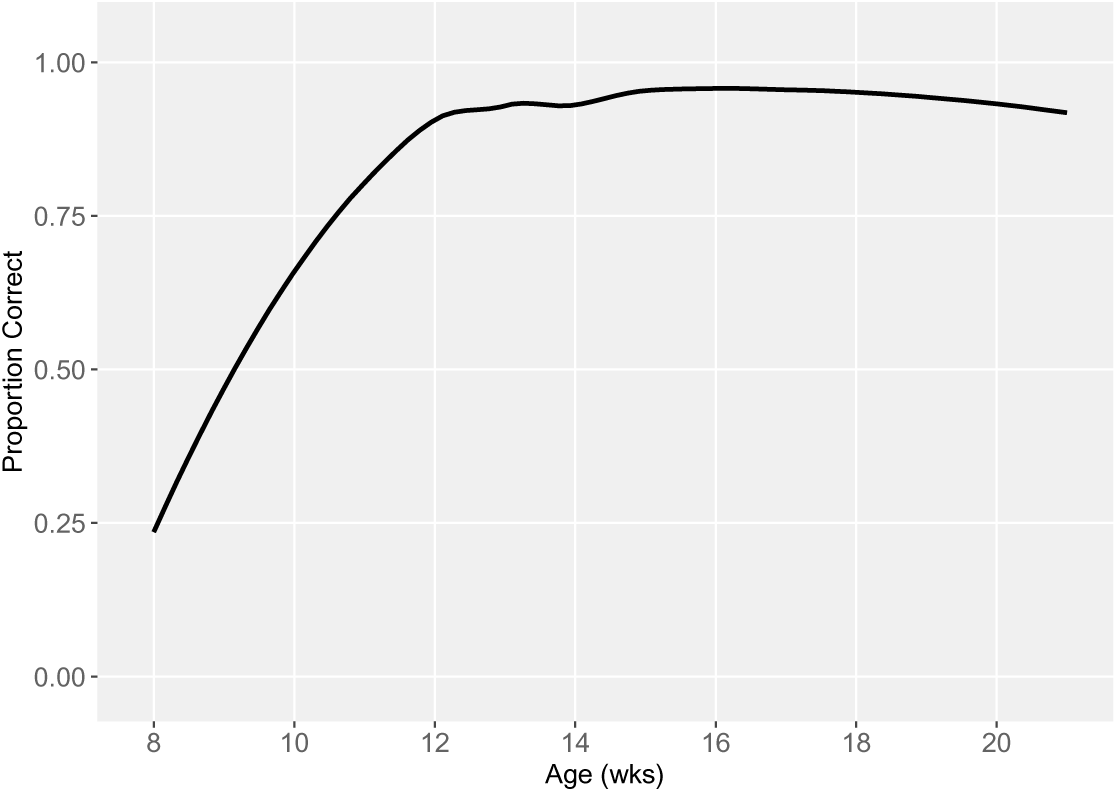
Performance on the Inhibitory Control trials of the Cylinder Task by Age in Weeks

#### Reversal Learning trials

Age (in weeks), rearing strategy (DPK vs home-raised), and experience (number of prior test sessions completed) were considered as covariates for generalized linear mixed effects models (**Table S8**). Age and experience were both significant predictor variables in all the models in which they were entered. Rearing was not a significant predictor variable in any of the models. The model with the lowest AIC score (1363.344) contained age and experience (β_age_= 0.057, SE = 0.026, p<.01; β_experience_ = 0.209, SE = 0.026, p<.001).

A two-tailed Welch’s t-test comparing the proportion of trials in which a clear route was taken to an open side without touching the transparent cylinder wall by the longitudinal subjects on their last testing session (only those aged 17-21 weeks old, *n =* 75, M ± SD = 0.62 ± 0.49) to that of control subjects in the same age range (*n =* 20, M ± SD = 0.28 ± 0.45) shows that the longitudinal puppies performed significantly better than the control puppies (t(133.18) = -5.98, p<.001) (Table 3, **Figure S19**).

Figure 8 and **Figure S20** graph all puppies’ performance on the inhibitory control reversal task across the 8-21 week age window using the “geom_smooth” function, method = ‘loess’ (Wickham, 2016), and **Figure S21** shows the same grouped by rearing strategy.

**Figure 8:**
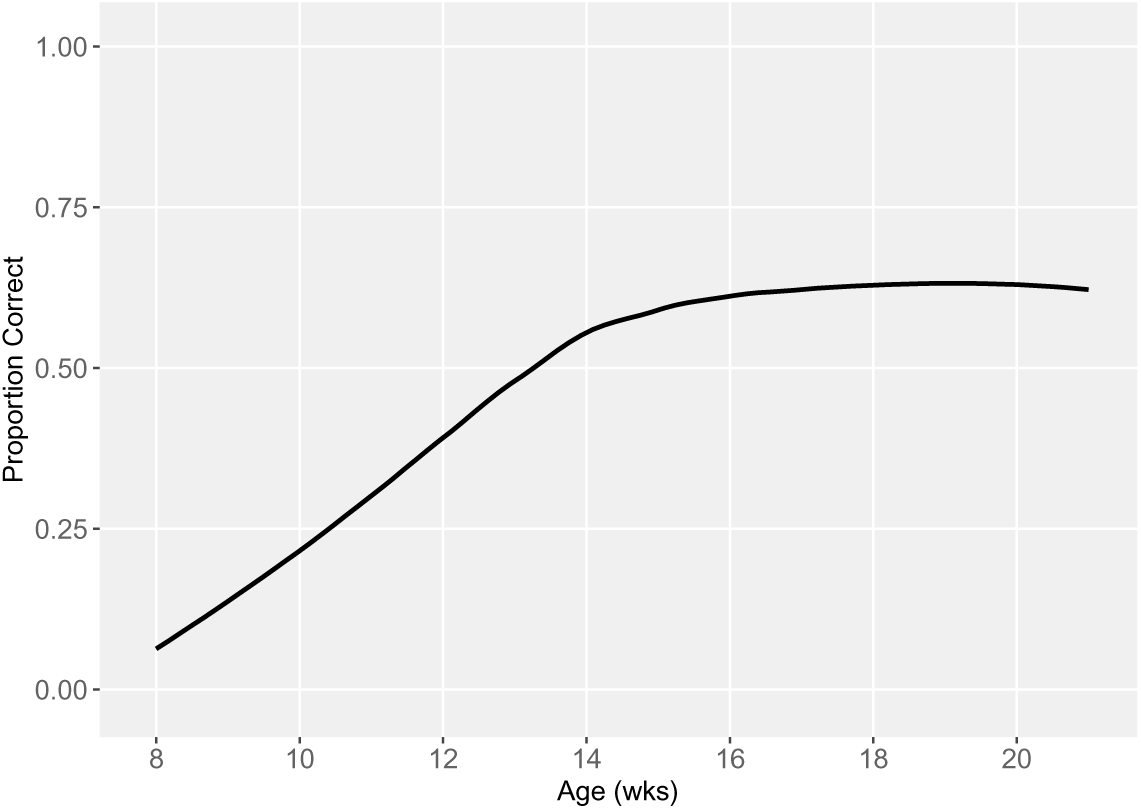
Performance on the Reversal learning trials of the cylinder task by Age in Weeks

#### Unsolvable (Eye Contact) Task

##### Total Looking Time

Age (in weeks), rearing strategy (DPK vs home-raised), and experience (number of prior test sessions completed) were considered as covariates for generalized linear mixed effects models (**Table S9**). Age and experience were both significant predictor variables in all the models in which they were entered. Rearing was not a significant predictor variable in any of the models. The model with the lowest AIC score (1,920.085) contained age and experience (β_age_= 0.194, SE = 0.075, p<.001; β_experience_ = 0.550, SE = 0.152, p<.001).

A two-tailed Welch’s t-test comparing the looking times (seconds) of the longitudinal subjects on their last testing session (only those aged 17-21 weeks old, *n =* 74, M ± SD = 5.34 ± 6.50) to that of control subjects in the same age range (*n =* 20, M ± SD = 3.07 ± 4.46) shows that the longitudinal puppies made significantly longer eye contact than the controls (t(184.91) = - 3.59, p<.001) (Table 3, **Figure S22**).

Figure 9 and **Figure S23** graph all longitudinal puppies’ (*n =* 91) mean looking times across the 8-21 week age window using the “geom_smooth” function, method = ‘loess’ (Wickham, 2016), and **Figure S24** shows the same grouped by rearing strategy.

**Figure 9:**
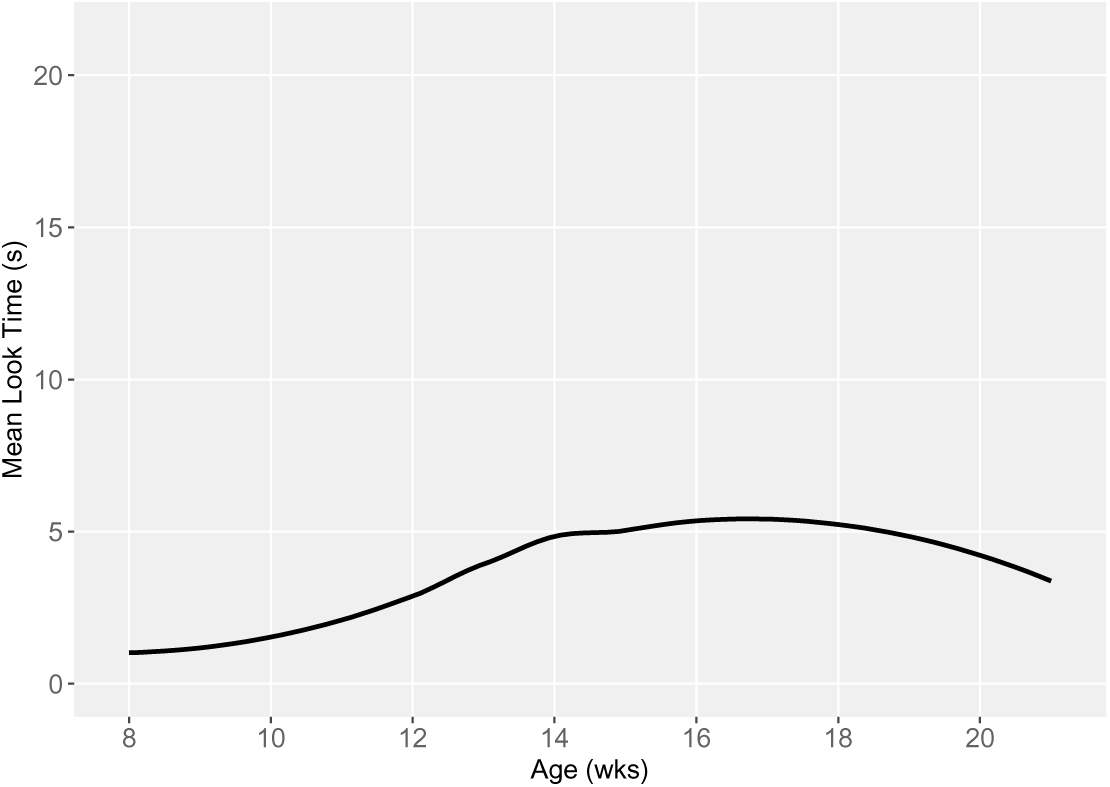
Performance on the Unsolvable Task (Looking Time) by Age in Weeks

##### Proportion of Trials Looked

Age (in weeks), rearing strategy (DPK vs home-raised), and experience (number of prior test sessions completed) were considered as covariates for generalized linear mixed effects models (**Table S10**). Age and experience were both significant predictor variables in all the models in which they were entered. Rearing was not a significant predictor variable in any of the models. The model with the lowest AIC score (1431.142) contained age and experience as predictor variables (β_age_= 0.081, SE = 0.031, p<.01; β_experience_ = 0.234, SE = 0.063, p<.001).

A two-tailed Welch’s t-test comparing the proportion of trials in which the longitudinal subjects made any eye contact with the experimenter during their last testing session (only those aged 17-21 weeks old, *n =* 74, M ± SD = 0.81 ± 0.39) to that of control subjects in the same age range (*n =* 20, M ± SD = 0.60 ± 0.49) shows that the longitudinal puppies made eye contact in a significantly higher proportion of trials than control puppies (t(109.44) = -3.47, p = <.001) (Table 3, Figure S25).

Figure 10 and **Figure S26** graph all longitudinal puppies’ (*n =* 91) proportion of trials in which they made eye contact across the 8-21 week age window using the “geom_smooth” function, method = ‘loess’ (Wickham, 2016), and **Figure S27** shows the same grouped by rearing strategy.

**Figure 10:**
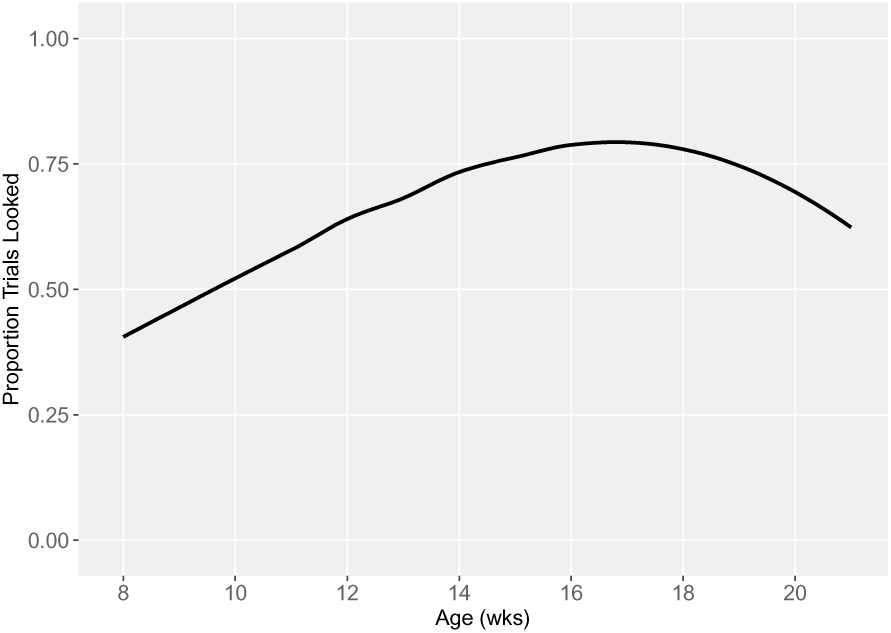
Performance on the Unsolvable Task (Proportion of Trials Looked) by Age in Weeks

### Causal Reasoning Task

Age (in weeks), rearing strategy (DPK vs home-raised), and experience (number of prior test sessions completed) were considered as covariates for generalized linear mixed effects models (**Table S11**). Age was a significant predictor variable in the age alone model and the age + rearing model. Experience was a significant predictor in all the models in which it was entered. Rearing was significant predictor variable in all the models in which it was entered. The model with the lowest AIC score (911.960) contained age, experience, and rearing as predictor variables (β_age_= 0.004, SE = 0.024, ns; β_rearing_= -0.354, SE = 0.143, p <.01; β_experience_ = 0.472, SE = 0.052, p<.001).

A two-tailed Welch’s t-test comparing the proportion of trials in which the baited bowl was chosen by the longitudinal subjects on their last testing session (only those aged 17-21 weeks old, *n =* 54, M ± SD = 0.89 ± 0.31) to that of control subjects in the same age range (*n =* 14, M ± SD = 0.43 ± 0.50) shows that the longitudinal puppies performed significantly better than the control puppies (t(59.68) = -6.25, p<.001) (**Table 3**, **Figure S28**).

Figure 11 and **Figure S29** graph all longitudinal puppies’ (*n =* 91) performance on the causal reasoning task across the 8-21 week age window using the “geom_smooth” function, method = ‘loess’ (Wickham, 2016), and **Figure S30** shows the same grouped by rearing strategy (intensive *n =* 35; extensive *n =* 77).

**Figure 11:**
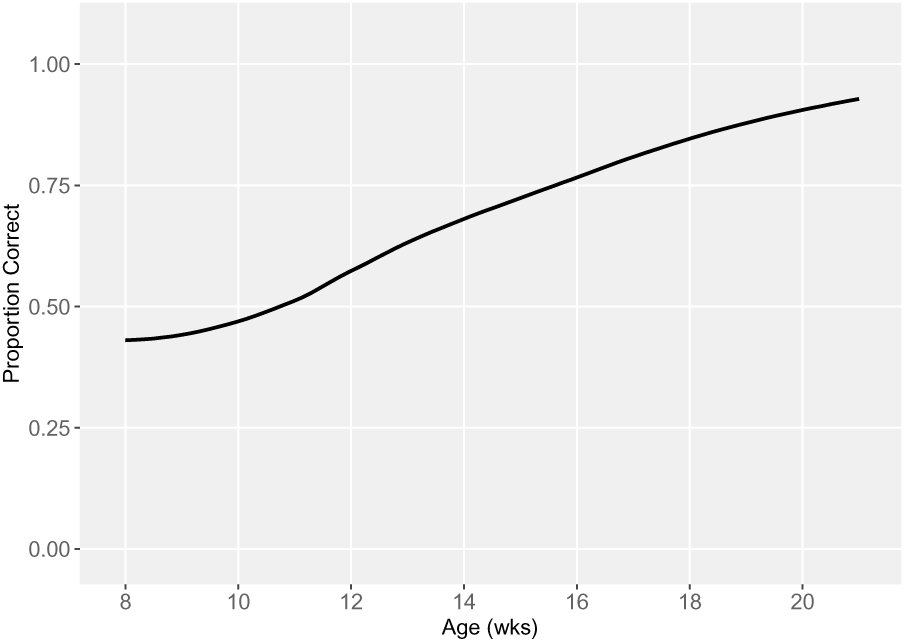
Performance on the Causal Reasoning Task by Age in Weeks

### Momentary Pointing Task

Age (in weeks), rearing strategy (DPK vs home-raised), and experience (number of prior test sessions completed) were considered as covariates for generalized linear mixed effects models (**Table S12**). None of these covariates were significant predictors in any of the models. The model with the lowest AIC score (1268.422) contained only rearing strategy as a predictor variable (β_rearing_=0.075, SE = 0.098, ns).

A two-tailed Welch’s t-test comparing the proportion of trials in which the baited bowl was chosen by the longitudinal subjects on their last testing session (only those aged 17-21 weeks old, *n =* 75, M ± SD = 0.51 ± 0.50) to that of control subjects in the same age range (*n =* 20, M ± SD = 0.48 ± 0.50) did not detect a difference in their performance (t(117.7) = -0.45, p=.65) (Table 3, Figure S31).

Figure 12 and **Figure S32** graph all longitudinal puppies’ (*n =* 91) performance on the momentary pointing gesture task across the 8-21 week age window using the “geom_smooth” function, method = ‘loess’ (Wickham, 2016), and **Figure S33** shows the same grouped by rearing strategy (extensive *n =* 35; intensive *n =* 77).

**Figure 12:**
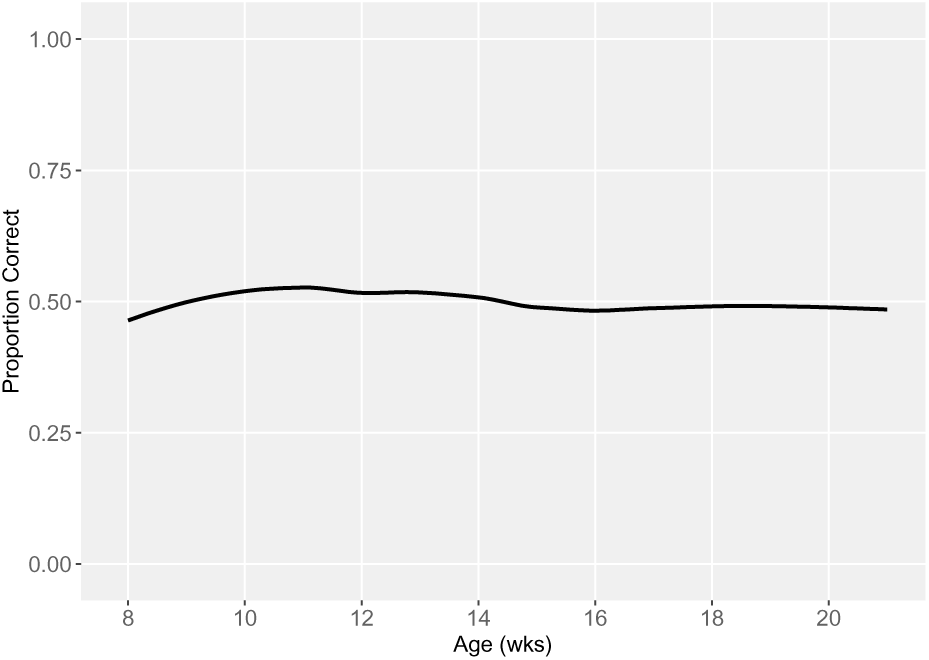
Performance on the Momentary Pointing Task by Age in Weeks

### Order of Rapid Development Completion (Breakpoint Analysis)

Of the skills which were fitted with segmented regression lines and breakpoints (see Methods), four (Marker Gesture, Cylinder Inhibitory Control, Cylinder Reversal Learning, and Unsolvable [Eye Contact]) were found to have a pattern of rapidly increasing performance, with a breakpoint where performance leveled out during this 8-21 week age window. One skill (Causal Reasoning) did not show this pattern, as performance instead continued to increase steadily across the age window. As such, the only breakpoint determined was very early, after a short period of decrease before the rapid increase began, and is likely an artifact of the relatively small sample size at the very young end of the age window (∼8 weeks). Both of the Working Memory skills (Delay and Distraction) showed slow steady increase across most of the age window, with a breakpoint after which performance seems to increase more rapidly towards the older end of the window. However, this could be due to a relatively small sample size at the older end (∼20 weeks) as well.

The four skills for which development reached a plateau during the tested age window are shown together (Figure 13), to compare the ages at which the performance on these skills began to level out, indicating the end of rapid development. The piecewise linear regressions and breakpoints for each of these skills are reported and graphed individually in the supplemental materials (Figures S34-S41).

**Figure 13:**
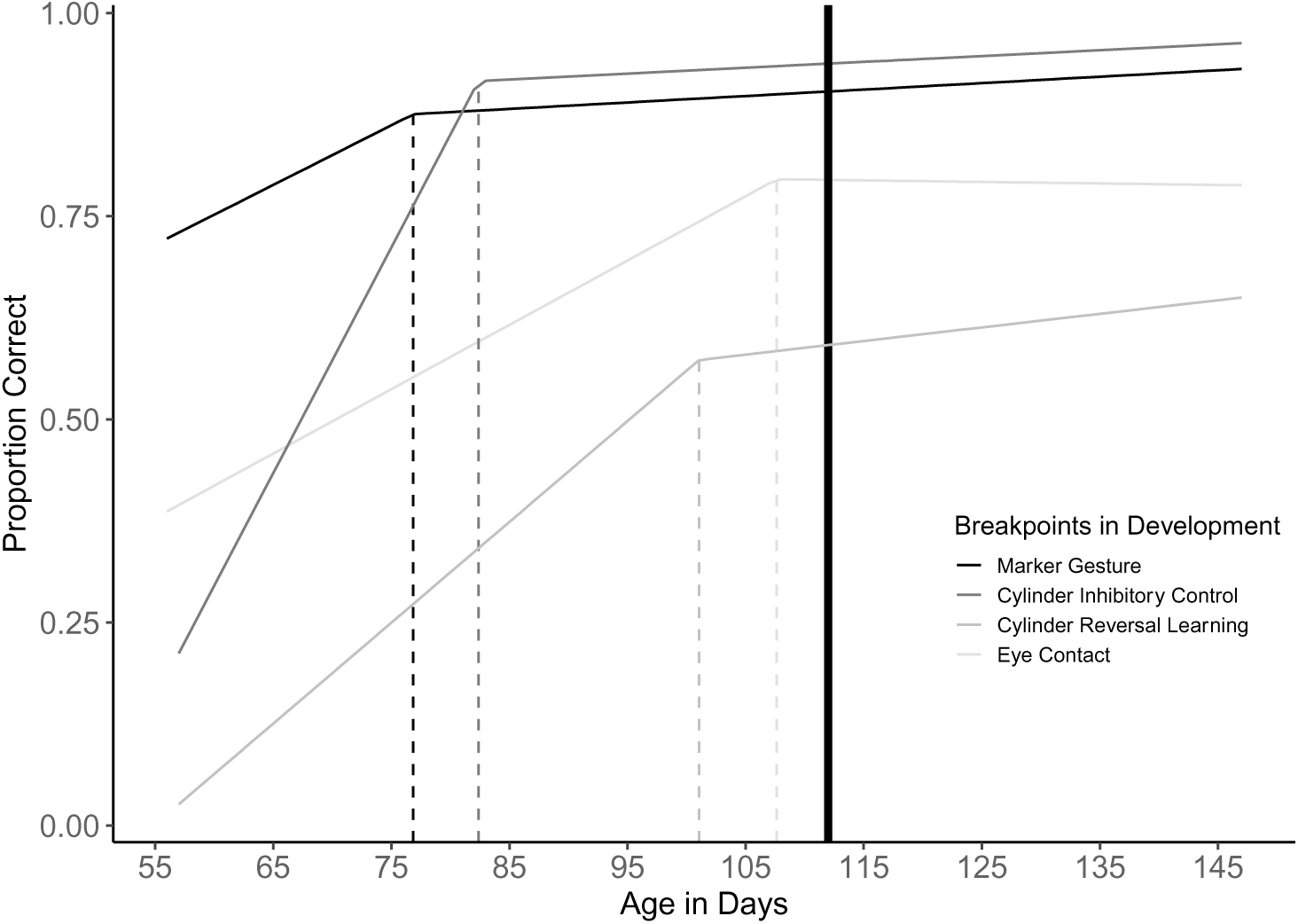
Combined plot of segmented regression lines for the four tasks which exhibit the pattern of rapid development completion within the 8-21 week age window. The solid line at 112 days represents 16 weeks or the end of the most rapid brain growth in puppies.

## General Discussion

The findings here provide strong support for the multiple intelligences hypothesis (MIH). Nine out of the ten skills measured were observed to emerge as puppies are completing a final stage of rapid brain growth before sixteen weeks of age. The pattern of emergence was not uniform across cognitive tasks nor tightly linked to previous experiences as predicted by either a unidimensional learning or general intelligence mechanism. Rather, cognitive skills emerged at different time points and developed at different rates while being differentially impacted by experience (**Table 4**). Development took a variety of paths, with skills developing in patterns ranging from rapid (inhibitory control) to gradual (causal reasoning). Developmental patterns differed between tasks, even though each task had the same number of trials (4), similar rewards, and were tested on the same schedule. Differential rearing had almost no impact on cognitive ontogeny. Extensive exposure to same-aged puppies and novel people and places on a college campus did not alter the development trajectory of social cognition in comparison to puppies intensively reared as part of a human family. Only a single non-social measure (causal reasoning) was impacted by rearing, with no ready explanation for this relationship. Repeated testing of the longitudinal puppies did not consistently affect their performance across tasks when compared to same-aged controls who were only tested once. While longitudinal dogs outperformed controls on some measures they performed similarly or less skillfully on others. The overall pattern is most consistent with the MIH. Different cognitive abilities emerge and develop at different ages and rates but show a highly similar maturation pattern even when puppies are intentionally raised two different ways. Experience impacts development, but across different cognitive abilities, maturational patterns vary in their plasticity, with some skills being impacted more by repeated testing than others (**Table 5**).

**Table 4.** Order of Emergence of Cognitive Skills. Performance on each task during bi-weekly testing. Dark grey shading indicates that the puppies, as a group, met emergence criteria (>50% of trials correct) for the skill at the significance level indicated by the stars:***p<.001, **p<.01, *p<0.5. “ns” indicates no significant difference from 50% correct, while significance stars with a < symbol indicate that performance was significantly below 50% correct. Note that on all two-bowl-choice tasks, the 50% emergence criteria represents chance, while on the cylinder and unsolvable tasks, chance cannot be defined and the criteria simply represents when as a group, puppies show the skill on more than half the trials.

|  | 8-9<br>Weeks | 10-11<br>Weeks | 12-13<br>Weeks | 14 – 15<br>Weeks | 16-17<br>Weeks | 18-20<br>Weeks |
| --- | --- | --- | --- | --- | --- | --- |
| Marker Gesture | *** | *** | *** | *** | *** | *** |
| Working Memory<br>(Delay) | *** | *** | *** | *** | *** | *** |
| Working Memory<br>(Distraction) | *** | *** | *** | *** | *** | *** |
| Auditory<br>Discrimination | *** | *** | *** | *** | *** | *** |
| Odor Discrimination | ** | ** | ns | ns | ns | ns |
| Cylinder (Inhibitory<br>Control) | ** < | *** | *** | *** | *** | *** |
| Unsolvable<br>(Proportion Looked) | ns | * | *** | *** | *** | *** |
| Causal Reasoning | ns | ns | *** | *** | *** | *** |
| Cylinder (Reversal<br>Learning) | *** < | *** < | * < | * | ** | ** |
| Momentary Pointing | ns | ns | ns | ns | ns | ns |

**Table 4:**
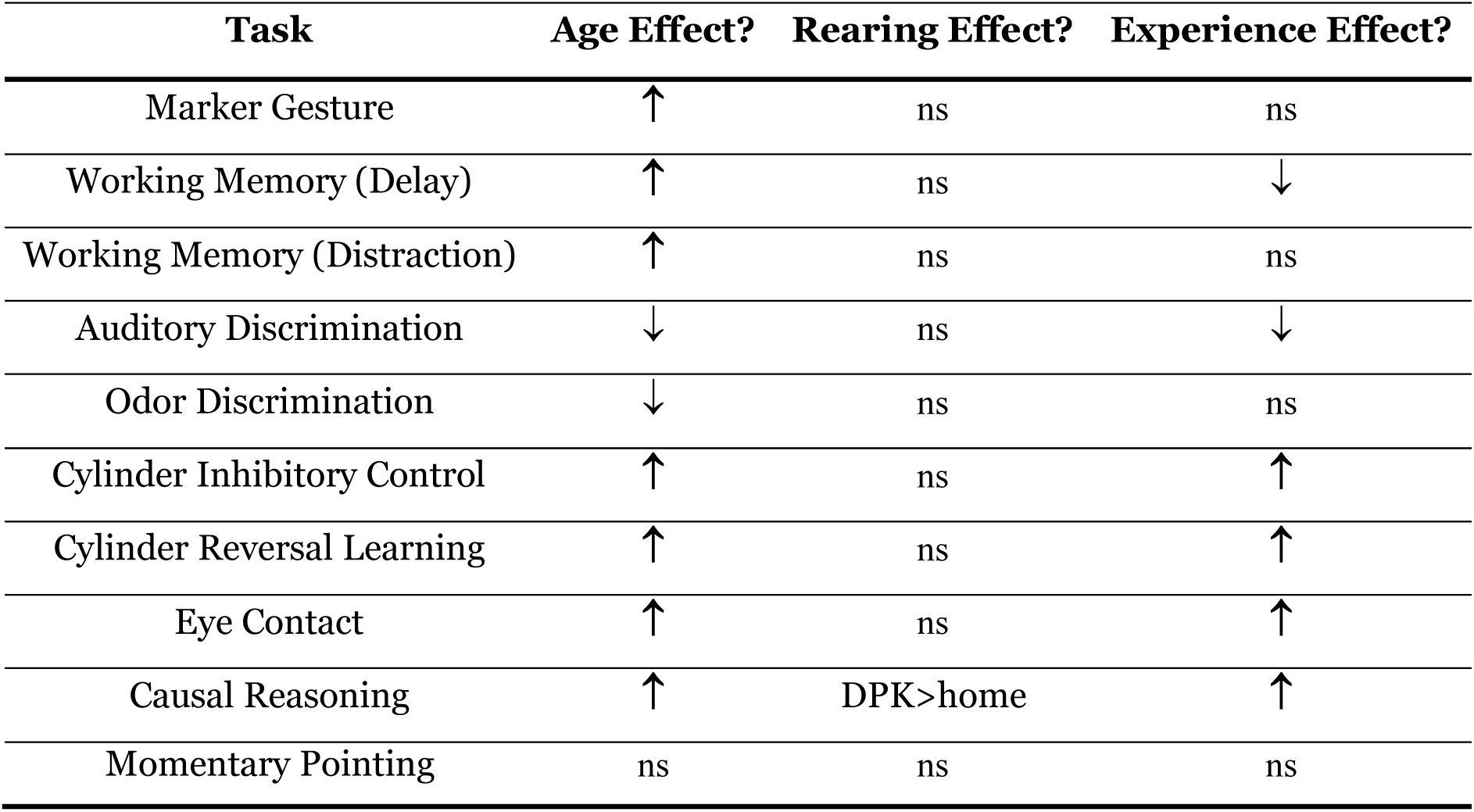
Summary of age, rearing, and experience effects on DCLB cognitive tasks. “↑” indicates a significant positive effect on performance, “↓” indicates a significant negative effect on performance, “ns” indicates no significant effect on performance.

The DCLB provides the highest resolution description of dog cognitive development currently available. The subjects all came from closed breeding populations that have been bred for assistance work for decades. All subjects were raised similarly prior to eight weeks of age and were tested using the same procedures (with high reliability observed across all measures). Despite these similarities, the DCLB revealed developmental variability in the emergence of cognitive skills. By defining ‘emergence’ as the longitudinal subjects’ group-level performance exceeding 50% of trials correct, we observe the following patterns (**Table 4**): First, the use of a novel gesture (marker), working memory, and sensory discrimination are present at 8-9 weeks, inhibitory control and eye contact (proportion of trials looked) emerge at 10-11 weeks, causal reasoning emerges at 12-13 weeks, and reversal learning emerges at 14-15 weeks. Only momentary pointing fails to emerge during the developmental window we examine (**Table 4**).

Second, testing for whether puppies reach a performance maximum followed by a plateau demonstrates how rates of emergence differed. For the marker gesture, inhibitory control, reversal learning and eye contact, the breakpoint analysis found the predicted change from a positive to near-zero slope, with each occurring at a different time before 16 weeks of age (Figure 13). Supporting the idea that these represent maturational changes, a subject’s age is also a significant explanatory variable in the model comparisons, with a positive beta coefficient in the best-fit model for all four of these tasks. In contrast, working memory and causal reasoning emerge in a different pattern. Both skills show steady development across test sessions with the best-fit models all including age as a predictor variable. However, puppies do not reach a ceiling or plateau within the developmental window tested. The observed developmental trajectory only leads to gradual improvement that likely continues after our final test period at 18-20 weeks. Meanwhile, both the sensory tasks showed regression instead of improvement over test sessions. The best-fit model for odor only contains age, while that for auditory discrimination only contains experience, with both having a significant negative beta coefficient on these respective variables. This drop in performance may have occurred as dogs became more reliant on visual information as their altricial visual cortex matured, and/or become expectant of visual cues as they gain experience with the other tests. The final pattern observed is a lack of development, with puppies showing no ability to use a momentary pointing gesture throughout the developmental window examined.

The DCLB also helps reveal what types of experience impact cognitive development and how that experience differentially impacts performance on the various tasks. First, the two rearing strategies had limited impact on cognitive emergence or the speed of development. The approaches were largely designed based on strategies that different assistance dog groups employ and differed on social variables previously shown to impact adult social behavior (e.g. Harvey et al., 2016; Vaterlaws-Whiteside & Hartmann, 2017). Perhaps most surprising is the lack of difference in performance on social tasks, despite half the puppies being extensively exposed to new people and other similarly aged puppies. The GIH predicts that through a general learning mechanism greater exposure to social interactions will alter the maturational course of social skills, but instead, we found that only the non-social causal reasoning task showed a significant effect of rearing strategy on performance. Second, comparing control dogs to the longitudinal dogs at the final testing time point (17-21 weeks) provides a powerful test of the impact of repeated testing on performance (**Table 3**). In six of ten tasks the control dogs performed as well or better than the longitudinal sample. In cases where longitudinal puppies were outperformed, it may have been that repeatedly completing a task they had succeeded with so early in life led to a loss of motivation and performance. For the remaining four tasks, the longitudinal dogs outperformed the same-age control dogs, making more eye contact in a higher proportion of trials and showing more inhibition, greater ability to reverse their learning, and more skill in the causal reasoning task. Repeated testing from an early age likely gave the longitudinal dogs an advantage, but the control dogs did perform above chance or have adult-like performance in the eye contact and inhibition tasks (see **Table S13**). This pattern suggests that when repeated testing positively impacted performance it did not cause a skill to emerge but instead interacted with brain maturation to enhance performance. It is also difficult to use reward history to explain the cases where the longitudinal dogs outperformed the controls because the unsolvable and cylinder tasks used to measure eye contact, inhibition, and reversal learning were all non-differentially rewarded. Only the causal reasoning task was differentially rewarded, but control puppies performed as well or better than longitudinal puppies on all the other object-choice tasks in which only correct choices were rewarded. It is easiest to interpret this pattern as a signal of maturation interacting with experience.

As a first attempt to closely examine the development of dog cognition during this final period of rapid brain development, our experiment has limitations. While we see significant development, in their first test session at 8-9 weeks of age, the longitudinal puppies performed above chance with half of our tasks (memory, marker and sensory tasks; replicating previous findings, Bray et al., 2020a; Bray, Gruen, et al., 2021; Salomons et al., 2021). This suggests that cognitive development occurring during the final period of rapid brain development tested here is a continuation of development occurring post-partum during nursing. Future research will be needed to examine the initial development of these skills before weaning (e.g. Riedel et al., 2008). Likewise, even in the final test session (i.e., typically after twenty differentially rewarded trials across six biweekly sessions), longitudinal puppies failed to comprehend momentary pointing. Adult dogs can read this gesture (Salomons et al., 2024), and puppies from the same assistance dog population can read the same gesture when the point is held statically as they make their choice (Bray et al., 2020), raising the question of when the capability to use more subtle communicative gestures first emerges. The DCLB is also largely designed based on tasks associated with or even predictive of adult training outcomes in working dogs (e.g. Bray et al., 2019; Bray, Otto, et al., 2021; Bray, Sammel, Cheney, et al., 2017; MacLean & Hare, 2018). Future research can be designed to examine the development of a larger range of social and nonsocial cognitive abilities. Particularly interesting might be the set of tasks previously shown to have a more human-like pattern in adult dogs than in great apes(MacLean, Herrmann, et al., 2017). As assistance dogs, our subjects are also only sampled from one subpopulation of dogs. These dogs are known to differ from pet dogs in having xenophilic responses toward strange people, and higher circulating baseline concentrations of oxytocin (Bray et al., 2019; MacLean, Gesquiere, et al., 2017). Many of the developmental patterns observed here may be specific to this population. A future priority will be to use the DCLB with a range of other dog populations to test for generalizability.

While recognizing the need for further research, our current findings have important implications for a puppy’s potential to learn during development. Understanding when different cognitive abilities emerge informs when different types of skills are most likely to be successfully learned. Puppies will likely succeed when tasks requiring self-control are introduced after 12 weeks of age, or even 14 weeks of age for those tasks requiring the most self-control (such as reversal learning). Recognizing the different speeds with which different abilities develop also hints at how quickly puppies might learn different skills. Tasks that require self-control may rapidly increase after 12 weeks of age while problems recruiting working memory may only gradually improve. Evaluating the impact of experience also suggests ways that social interactions may or may not enhance social learning in puppies. While our different rearing strategies produced no cognitive differences, participating in the unsolvable task for a few minutes every other week almost doubled the amount of eye contact longitudinal puppies made in this task. This type of task might be used to increase eye contact and social bonding between humans and dogs. Future research can examine if early exposure to the unsolvable tasks leads to more eye contact in other contexts as dogs reach adulthood. There is also the potential to re-test our subjects on the DCLB when they reach adulthood, to further examine the stability of cognitive performance across the life span. Critically, this will include testing whether individual differences in cognitive performance during puppies’ development are related to their professional training outcomes as adults. It may be that evaluations of each cognitive skill conducted soon after that skill has emerged may lead to the strongest predictions, which can inform the timing of cognitive evaluations of assistance dog puppies in the future. Previous work also suggests that combining cognitive and temperament measures provides increased predictive power of adult training success (Bray, Sammel, Cheney, et al., 2017; Bray, Sammel, Seyfarth, et al., 2017; Lazarowski et al., 2019; Smith et al., 2024). Combining developmental measures of temperament with our cognitive measures may again provide the most powerful way to anticipate training outcomes before an adult dog is even enters professional training.

Taken together, and despite the limitations, the results further support prior research demonstrating different cognitive domains in dogs. The cognitive abilities evaluated by the DCLB develop separately, at different rates, and during a final period of rapid brain growth in puppies. Cognitive skills do not uniformly improve in synchrony with age and different types of experience as predicted by a unidimensional, generalized mechanism. Age was repeatedly the defining factor explaining patterns of performance, but the comparison of the longitudinal and control dogs shows that experience plays an important role in some cases as well. Based on our findings, the question is not whether cognition and the developing brain is impacted by experience, but rather, to what degree and by which types of experience? Some types of cognition appear to be more plastic than others: for example, the amount of eye contact made during the unsolvable task is much more affected by previous experience than is the ability to read human gestures (**Table 5**). Finally, this type of research with dogs also has the potential to provide a roadmap for testing how cognitive architecture develops and evolved in a wider range of species.

## Acknowledgements

Thanks to all the wonderful staff and volunteers at Canine Companions, Guiding Eyes for the Blind, and Eyes, Ears, Nose and Paws, who made this collaborative effort possible. Thanks to Philip White for assistance with analysis. Thanks to the many research assistants who helped with puppy handling and data entry/coding, especially Gabrielle Bunnell, Charlotte Burnham, Samantha Dunn, Samantha Kefer, Emma Nelson, Jessica Shoemaker, Jordan Sokoloff, Jamie Sokoloff, Elizabeth Wise, Jacqueline Wright.

## Ethics Approval

This work was conducted under Duke University IACUC protocol A122-21-06.

