## Supplementary Material for "Longitudinal evidence for the emergence of multiple intelligences in assistance dog puppies"

### Supplemental Methods

#### Dog Cognitive Longitudinal Battery (DCLB)

| Domain | Subdomain | Test | Associated w graduation?[1] | Sig. development 10 wks -> adult?[2] | Consistent w/in individuals 10 wks -> adult?[2] |
|---|---|---|---|---|---|
| Non-Social Cognition | Memory | Warm-Ups (Spatial Memory) | ✓ | ✗ | ✗ |
| | | Working memory (20 sec delay) | ✓ | ✓ | ✗ |
| | | Working memory (distraction) | ✓ | ✓ | ✗ |
| | Physical Reasoning | Causal reasoning | ✓ | (not tested) | (not tested) |
| | Sensory Perception | Auditory discrimination | ✗ | ✓ | ✗ |
| | | Odor discrimination | ✓ | ✓ | ✗ |
| | Inhibition | Cylinder | ✓ | ✓ | ✓ |
| Social Cognition | Communicative Intentions | Momentary pointing | (not tested) | (not tested) | (not tested) |
| | | Marker gesture | ✓ | ✓ | ✓ |
| | Referential Communication | Unsolvable task (eye contact) | ✓ | ✓ | ✗ |

*Table S1: The DCLB consists of ten cognitive tasks validated for use in puppies and adult dogs in previous work. Subdomains refer to the cognitive abilities each test is designed to assess. Check marks indicate where a test was previously associated with working dog training success, showed development over time, and/or was stable in dogs tested as puppies and then again as adults. An X indicates where a test did not meet one of these criteria (based on 1) MacLean & Hare, 2018; 2) Bray et al., 2020).*

#### Task-Specific Method Details

*Choice Criteria for Object-Choice Tasks*

*This included the Marker Gesture, Working Memory, Momentary Pointing tasks.* A choice is defined as the subject's snout or front paw crossing the rim of a bowl. When a choice is made, H says, "choice". If E sees the choice first, despite looking down, E may also say "choice". If the subject chooses the baited bowl, E praises the subject and allows them to eat the reward. If the subject chooses the incorrect bowl, E says "wrong" in a neutral, monotone voice. E then returns the subject to H, retrieves the reward if the subject chose incorrectly, and the next trial can commence.

*Bowl Choice Warm-Ups*

*Setup and Familiarization (Phase 1):* H holds subject in the starting box. E kneels behind the experimenter's line. E presents the reward and then visibly places the food in the bowl 1) halfway between the puppy and experimenter and 2) then directly in front of E on the 1-m testing line. On each trial the puppy is released to obtain the reward. First, E shows food to the puppy in hand and says "Puppy, look!", then when E places the food in the bowl, she says "Puppy, look!" again. E rests with her hands behind her back, looks straight down, and says "okay". The subject is then allowed to approach the food and obtain the reward. If the subject does not approach the kibble in 20 s, the trial is repeated. If the puppy approaches/climbs on E or sits and waits for E, the trial may be repeated until the puppy retrieves the kibble immediately on its own. After retrieving the food successfully from each location one time, the subject moves on to Phase 2.

*One Bowl Alternating (Phase 2):* E presents the reward and visibly baits a single bowl placed in one of the two positions on either side of E's position (i.e. the 10-cm circles at either end of the 1-m test line; see Figure 1). After baiting, E kneels at the E location in resting position. The subject is then allowed to approach the bowl and obtain the reward. If the subject does not touch/choose the bowl in 20 s, the trial is repeated. This phase of warm-ups introduces subjects to the two locations they can find rewards and assures that the subject is motivated to find them. Returning to these trial types during testing can serve as a correction procedure for spontaneous side biases (i.e. a subject repeatedly chooses one side only) and ensures that subjects gain experience finding the reward in both locations. The subject is required to successfully retrieve the reward on four trials, twice on each side, to move on.

*Two Bowl (Phase 3):* Two bowls are simultaneously placed in the 10-cm circles connected to the 1-m line (Figure 1). E presents the reward to the puppy and visibly baits one of the two bowls according to the predetermined order on the data sheet. After baiting, E kneels in resting position. The subject is then allowed 20 s to make a choice. If the subject chooses the baited bowl, the subject is allowed to have the reward, E praises the subject, and the next trial is administered. If the subject chooses the incorrect bowl,

E says, "Wrong" in a neutral tone and the subject is not rewarded. If the subject does not choose any bowl ("no choice") within 20 s, the trial is repeated. This phase of warm-ups assures that the subject is not choosing bowls randomly and is fixing attention on the experimenter's actions. Subjects should choose correctly on their first attempt in four out of five consecutive trials to advance to test trials within a maximum of 12 trials (including repeated trials, but not refamiliarizations). If subjects do not pass this phase within that number of trials on the last possible attempt of warm-ups, they may continue to test trials if they have been consistently choosing a bowl, but not if they have not passed due to no-choice responses.

*Refamiliarization/Abort Criterion:* If the subject does not choose by touching a bowl within 20 seconds on two trials in a row during phase 2, phase 1 is resumed until the puppy reengages; then phase 2 trials are resumed. If the subject does not choose on two trials in a row during phase 3, two trials of phase 2 are conducted (one-bowl alternating). If there are 4 no choices (NCs) total in either phase 2 or phase 3 (including refamiliarizations), the session is aborted.

<u>*Marker Gesture*</u>

*Procedure:* E places both bowls next to each other at location C. The occluder is placed ~10 cm in front of the bowls, blocking the puppy's view of the bowls. E reaches over the occluder to present the subject with the reward (saying "Puppy, look!"), then places the treat in one of the bowls behind the occluder while simultaneously sham baiting the other bowl with the other hand (i.e. mimicking the baiting movement with the empty hand). E then places the occluder behind her and simultaneously slides the bowls into locations R and L. As E finishes sliding the bowls into the circles marked on the mat at R and L, she glances first to the bowl on her left, and then to the bowl on her right to check placement. E spends equal time glancing at each bowl placement and always in left-right sequence. Once the bowls are in place, E picks up the yellow marker from the center of the mat and using her right hand, leans forward to show it to the puppy (~10 cm from the subject's nose), and says "puppy, look!" while attempting eye contact. E then says "puppy, look!" again while placing the marker next to the appropriate bowl (in the square marked on the mat, see Figure 1) and directing gaze towards the marker/baited bowl. E then returns to the resting position and gives an “okay!” release, signaling H to release the subject to make a choice. Until this point, H should be holding the puppy but looking down so that H is blind to which side is being indicated. Once E has given the “okay!” release signal and the puppy has left the starting box, H looks up; when the puppy makes a choice, H says, “choice”. E maintains resting position until the subject makes a choice or the maximum trial time of 20 s has elapsed. Four trials are conducted and the dependent measures is whether the puppy correctly chose the baited and marked bowl on each trial.

*Refamiliarization/Abort Criteria:* If the subject does not make a choice within 20s, the trial is repeated. If the puppy no-choices (NC) twice in a row, 2 trials of two-bowl alternating warm-ups are run. If the puppy does not engage in refamiliarization trials or makes 4 total no-choices, the test is aborted.

*Working Memory (Delay)*

*Procedure:* The puppy is positioned in the starting box. Two bowls are positioned in the 1-m bowl circles. H holds a countdown stopwatch set for 20 sec. E places a running stopwatch in count-up mode on their kneeling pad to count the 20 sec interval for each trial. E presents the subject with the reward, saying “Puppy, look!”, and then visibly places the reward into one bowl while saying “Puppy, look!” again. E then looks down at the running stopwatch and counts 20 seconds of delay. When the time is up, E says, “okay!” and H releases the puppy and starts the timer. The trial ends when the puppy makes a choice or after 20 s has elapsed. Four trials are conducted, and the dependent measure is whether the puppy correctly choses the baited bowl on each trial.

*Refamiliarization/Abort Criteria:* If the subject does not make a choice within 20s, the trial is repeated. If the puppy no-choices (NC) twice in a row, 2 trials of two-bowl alternating warm-ups are conducted before resuming. If the puppy does not engage in refamiliarization trials or makes 4 total no-choices, the test is aborted.

*Working Memory (Distraction)*

*Procedure:* E engages the puppy to play with a toy for 1 – 2 minutes. If the puppy seems uninterested in this toy, E switches to a toy that is of higher interest so that it can create a distraction during the trials. A large digital display clock is positioned on a shelf/table behind H such that it is visible to E to count the 20 sec duration for toy distraction for each trial. The puppy is positioned in the starting box. Two bowls are positioned in the 1-m line bowl circles. E presents the subject with the reward (until the distraction begins the toy is placed behind E's back so that it is out of the puppy's view), saying "Puppy, look!" and then visibly baits one bowl (saying "Puppy, look!" again). E then begins distracting the puppy (being held by H) with the toy, behind the sniff line (see **Error! Reference source not found.**) in front of the puppy. E engages the puppy by spinning the toy on the mat, squeaking it, talking to the puppy, etc. The puppy is allowed to touch and interact with the toy during the distraction time while staying behind the sniff line. E glances at the large display clock behind H and at 18 secs stops the distraction, places the toy behind her back, and sits in a resting position (20 sec total elapsed time). E then says, "okay!" and H releases the puppy and starts the timer. The trial ends when the puppy makes a choice or after 20s have elapsed. Four trials are conducted, and the dependent measure is whether the puppy correctly chooses the baited bowl on each trial.

*Refamiliarization/Abort Criteria:* If the subject does not make a choice within 20s, the trial is repeated. If the puppy no-choices (NC) twice in a row, 2 trials of two-bowl alternating warm-ups are completed before resuming. If the puppy does not engage in refamiliarization trials or makes 4 total no-choices, the test is aborted.

*Auditory Discrimination*

*Familiarization:* Familiarization consists of 2 trials. H holds the puppy in the starting box; E is kneeling at a central position, directly in front of the puppy, behind the 50 cm bowl circles. E places a bowl 50 cm from the starting box in the right circle (see Figure 1). E presents the subject with dry kibble, saying "puppy look!", and baits the bowl. When baiting, E drops the kibble in the very front of the bowl (so the puppy cannot see it) and from a height of 1-2 cm (so that it makes an audible "clink" cue). E says "Okay!", H releases puppy, and puppy is allowed to retrieve kibble from the bowl. Repeat with kibble dropped into the bowl placed in the left circle at 50-cm. The puppy has 20 sec to retrieve the reward.

*Test Procedure:* E places bowls in 50-cm circles. E presents subject with dry kibble, saying "puppy look!", baits and sham baits the bowls. When baiting, E drops the kibble in the very front of the bowl (so the puppy cannot see it) and from a height of 1-2 cm (so that it makes an audible "clink" against the metal bowl). When sham baiting, E puts her hand about 2 cm from the bottom of the bowl, pauses, and relaxes her grip/moves a finger slightly to imitate dropping. Bowls are always baited/sham baited in the order right then left. The subject is given 20s to make a choice in four test trials, and the dependent measures is whether the puppy correctly chose the baited bowl on each trial.

*Refamiliarization/Abort Criteria:* If the puppy does not choose within 20s on 2 consecutive trials, two familiarization trials are run to encourage them to eat from a bowl. If there are 4 total no-choices, the test is aborted.

*Odor Discrimination*

*Apparatus:* PVC elbows are set up with one designated as the odor elbow (labelled "food") and the other as the blank (labelled "clean"). 10 dry pieces of kibble are loaded into the odor elbow. Then, a cotton pom pom large enough to fill the opening is placed inside the elbow directly behind the cloth mesh, which has been permanently secured between the two PVC pieces to block visual access to the treats, but still allows odor to dissipate. The blank, or "clean", elbow is assembled in the same manner, but absent of treats. Two pom poms large enough to completely fill the openings are used at the top of each elbow to close the top and allow a treat to be placed on top of the odor elbow for fast delivery of a reward to the puppy. E holds the elbows from above with the palms covering the upper opening so that the puppy cannot access the treat or the top of the elbow. The longer end of the apparatus is the end that is presented to the puppy.

*Familiarization:* H positions the puppy in the starting box (S, **Figure 1**). E says "sniff it" in a high, encouraging tone and presents the puppy with the odor elbow 10 cm in front of the puppy box and at the height of the puppy's nose. After 3 seconds, E pulls the elbow back to 50 cm away and holds it in place on the mat, centered between the $R^1$ and $L^1$. The puppy is given 20s to touch the elbow and is rewarded immediately upon touching. The first trial the elbow should be held in the right hand and the second in the left hand. A trial is repeated if the puppy does not choose.

*Test Procedure:* H positions the puppy in the starting box. E kneels behind $R^1$ and $L^1$, so that she is within arms' reach of the puppy positioned in the start box. E puts both elbows next to each other and centered between $R^1$ and $L^1$ and then presents them simultaneously to the puppy at the front edge of the start box, at the height of the puppy's nose. The puppy is given 3s to smell them. E says "sniff it" to encourage the puppy to smell the elbows. E then pulls the elbows back, following the slanted lines on the mat and places the elbows on the ground at $R^1$ and $L^1$. E says, "okay!" and H releases the puppy to make a choice. The puppy must touch the elbow or the experimenter's hand (wrist, back of hands or fingers) to count as a choice. If the puppy chooses the baited elbow, the puppy is rewarded. The trial ends when the puppy makes a choice or after 20s has elapsed. Four trials are conducted and the dependent measure is whether the puppy correctly choses the baited elbow on each trial.

*Refamiliarization/Abort Criteria*: If the subject no-choices twice consecutively, two refamiliarization trials are conducted: the subject is presented with a single elbow and is rewarded for approaching. If the puppy does not approach on either refamiliarization trial, or if there are 4 total "no-choices", the test is aborted.

<u>*Cylinder (Inhibitory Control & Reversal Learning)*</u>

*Apparatus and Setup:* A transparent cylinder was constructed by cutting an acrylic aquarium (Koller Products 6 gallon AquaView 360) to a cylinder 1ft wide x 11in diameter. An opaque gray fabric cover was made to fit around the cylinder, which completely covered the transparent walls, and could be affixed to and removed from the cylinder with Velcro at the base. The cylinder was glued to a square wooden base (2ft x 2ft) to keep the apparatus still if bumped by the subject. E kneels centered behind the cylinder (E's knees may be on the board that cylinder is attached to), while H centers the puppy in the starting box. The cylinder should be centered at location C, with the longitudinal midline of the aligned with the bowl aligned between locations R and L.

*Warm-up:* E places kibble in bowl, shows puppy, says "Puppy, look!" and places the bowl about 50 cm from puppy. E says "okay" and allows the puppy to eat the kibble out of the bowl. The subject has 20 s to approach the bowl. After the puppy has successfully approached the bowl at 50 cm, the procedure is repeated with the bowl right in front of the cylinder (~90 cm from puppy). Once the puppy has approached the bowl and cylinder, leading trials were initiated. E shows the subject the bowl with the reward, H releases puppy when E says "okay!", and E uses the bowl to guide the puppy into the side of the tube. These warm-ups are repeated as needed until the puppy succeeds at following the bowl into the tube for two trials, one baited from the right side and one from the left. During any of these warm ups, if the subject did not approach and retrieve the reward, E worked to make the puppy comfortable (petting, encouragement, playtime if necessary) and repeated the procedure. If a subject failed to approach during these repeated trials, E increased the food reward. If a subject still did not approach on the next two trials, the test was aborted.

*Familiarization (opaque cylinder):* E says "Puppy, look" while showing the subject the bowl and reward, and baits the opaque cylinder with the right hand. E is in resting position and says "okay!". H then starts a 30-s timer and releases the puppy to make a choice.  Subjects are permitted (and encouraged) to retrieve the reward on all trials regardless of the accuracy of their first attempt or if they chose at all. In the case that the puppy touches the cylinder but never retrieves the reward on its own, E shows the puppy the solution and gives the puppy the reward. When possible, E removes the bowl from the side opposite the puppy while H retrieves the puppy. On each trial, E scores whether the subject made a choice without touching (1) or touched before choosing (0), and which side they chose (L/R from E's perspective). A choice consists of the puppy's snout or front paw crossing the plane at either open end of the cylinder. If the puppy brushes the edge of the cylinder in the process of going in the side, it is coded as a correct response, not a touch. If the puppy's ears, but nothing else, brush the cylinder at any point, this is also not considered a touch. Before advancing to the tests, subjects must correctly (score of 1) retrieve the reward in 4 out of 5 trials within a maximum of 12 familiarization trials.

*Inhibitory Control Trials (transparent cylinder):* The test trials are identical to familiarization trials except that the fabric cover is removed so the cylinder is transparent. As before, E codes whether the subject made a choice without touching (correct) or touched the transparent wall before choosing (incorrect), and which side they chose (L/R from E's perspective). Again, subjects are allowed to retrieve the reward on all trials regardless of the accuracy of their first attempt. Four trials are conducted.

*Reversal Learning Trials (transparent cylinder):* The reversal trials are identical to the test trials except that after the cylinder is baited and before the subject is called, E places a clear barrier over one side of the cylinder. The blocked side is determined by whichever side was chosen in the majority of the last three detour trials. This is also the hand with which E baits the cylinder for this phase. As before, the puppy has 30 s to make a choice and is rewarded regardless of accuracy (i.e. allowed to eat the treat when found even if previously touched cylinder). If the puppy touches the cylinder but never solves t`he problem on its own, the puppy is shown the solution after 30 seconds. To do so, the experimenter calls the puppy's attention to the treat. If that's not enough, the experimenter may gently guide the puppy toward the open side of the cylinder to get the treat. If the puppy does not make a choice within 30s, they are not rewarded and the trial is repeated. Four trials are conducted.

*Refamiliarization/Abort Criteria:* In familiarizations, if two consecutive no choices occur, two more leading trials are conducted before resuming. If the dog does not meet the criterion to pass familiarizations within 12 trials, the task is aborted. In detour and reversal trials, if two consecutive no choices occur, the food reward is increased. If the subject does not choose on a total of 4 trials across phases, the task is aborted.

*Unsolvable (Eye Contact)*

*Apparatus:* A clear container with a clear lid that snaps closed (Rubbermaid Brilliance 1.3 cup container or similar) is super-glued to a wooden base (2ft x 2ft) so that it cannot be picked up by the subject.

*Warm-up Trials (Solvable):* H centers the subject in the start box (S, Figure 1) and gently holds the subject in place. E kneels at location E, behind the open clear container with the lid off, which is positioned at location C. E reaches forward (over the container) to present the food reward and says “Puppy, look!”, allowing the subject to see and sniff the food briefly. E then places the food inside the container, and positions the lid loosely on top such that it could easily be knocked off. E then places her hands on her thighs, looks down, and says “Okay!”, signaling H to let go of the subject. The subject is allowed to knock off the lid and retrieve the reward, at which point E provides verbal praise and the trial ends. If at any point during the trial the subject somehow knocks the lid into a position which makes the reward unreachable, E quietly intervenes to make the reward accessible again. If the subject does not successfully retrieve the reward within 30s, the trial is repeated. The subject is required to successfully complete 4 of these warm-up trials before advancing to the test. The first of these starts with the lid leaning up against the container, the second with the lid covering about half of the opening of the container, and the third and fourth with the lid covering three-quarters of the opening of the container.

*Test Trials (Unsolvable):* Test trials are identical to warm up trials except that E seals the lid onto the container so that it cannot be removed by the subject. The subject is given 30 seconds to attempt to access the reward. E remains seated in place and continuously looks at the subject (rotating head and body if necessary) during this period. E holds a silent stopwatch, and uses the start/stop buttons in count-up mode to measure the total time that the subject looks at their face. After each test trial, E praises the subject, opens the container, and allows the subject to retrieve the reward. Four test trials are conducted, and the number of seconds spent looking at E's face for each trial was used as the measure for analysis.

*Abort Criteria:* The session is aborted if the subject does not complete all four familiarization trials within 4 attempts for each trial.

<u>*Causal Reasoning*</u>

*Familiarization:* E shows the puppy the reward inside of the bowl with the wooden block base attached, saying "Puppy, look!". Then the bowl is placed at location $L^1$ or $R^1$ (one familiarization trial done at each location). E sits in resting position, says "Okay!", and H releases puppy to retrieve the food. For the third familiarization test, the puppy is shown the reward in the same manner and then the bowl is placed at location C. An opaque yellow circular cloth (50cm diameter), held in both hands, is then draped over the bowl and the subject is allowed to approach. When they touch the cloth with their nose or front paw, E lifts the cloth and the puppy is allowed retrieve the food (in order to insure that subjects understand they can search and will get the food even if the bowl is covered). Once these three familiarization trials are completed, then the subject can move on to the test trials.

*Test Procedure:* Two of the yellow circular cloths are laid flat on the mat, one on each of the bowl locations. E shows the puppy the reward inside the bowl and then hides it behind the large occluder before beginning the hiding process. Moving left to right, at the location to be baited, E lifts the cloth above the occluder, places the bowl on the ground, and then places the cloth over the top of the bowl such that the bowl is covered by it but the shape of the bowl visibly displaces the cloth. At the unbaited location, E similarly lifts the cloth above the occluder and places it smoothly back on to the floor in the same location. The occluder is then removed and placed behind E. E sits in resting position, says "okay!" and H releases the puppy to make a choice. A choice is scored when the puppy touches its nose to the cloth. If the puppy choses correctly, then the cloth is immediately removed to allow puppy access to the reward. If the puppy makes the wrong choice, then the cloth is not lifted until the puppy returns to the starting point. The trial ends when the puppy makes a choice or after 20s has elapsed. Four trials are conducted and the dependent measure is whether the puppy correctly chooses the baited bowl under the raised cloth on each trial.

*Refamiliarization/Abort Criteria:* If the subject does not make a choice within 20s, the trial is repeated. If the puppy no-choices (NC) twice in a row, 2 trials of two-bowl alternating warm-ups are completed. If the puppy does not engage in refamiliarization trials or makes 4 total no-choices, the test is aborted.

<u>*Momentary Pointing Gesture*</u>

*Procedure:* E places both bowls next to each other at location C. The occluder is placed ~10 cm in front of the bowls, blocking the puppy's view of the bowls. E reaches over the occluder to present the subject with the reward (saying "Puppy, look!"), then places the treat in one of the bowls behind the occluder while simultaneously sham baiting the other bowl with the other hand (i.e. mimicking the baiting movement with the empty hand). E then places the occluder behind her and simultaneously slides the bowls into locations R and L. As E finishes sliding the bowls into the circles marked on the mat at R and L, she glances first to the bowl on her left, and then to the bowl on her right to check placement. E spends equal time glancing at each bowl placement and always in left-right sequence. After baiting, E tries to make eye contact with the puppy, says "puppy, look!", and points toward the bowl with the reward. The point should consist of the index finger of the contralateral hand extended about 20 cm from the bowl (other hand should already be behind E's back) and E's head and gaze directed toward the bowl . This point is held for 2 seconds ("1, Mississippi 2, Mississippi) and then E places hand behind back and sits in resting position. E then gives an "okay!" release signaling H to release the subject to make a choice. Until this point, H should be holding the puppy but looking down so that H is unaware which side is being indicated. Once E has given the "okay!" release signal and the puppy has left the starting box, H looks up; when the puppy makes a choice, H says, "choice". The maximum trial time is 20 s for the puppy to make a choice. If the puppy makes the incorrect choice, E says "wrong" in a neutral tone. The puppy is not given a reward or shown the correct bowl, but does advance to the next trial. Four trials are conducted and the dependent measure is whether the puppy correctly choses the baited and point-indicated bowl on each trial.

*Refamiliarization/Abort Criteria:* If the subject does not make a choice within 20 s, the trial is repeated. If the puppy no-choices (NC) twice in a row, 2 trials of two-bowl alternating warm-ups may be done. If the puppy does not engage in refamiliarization trials or makes 4 total no-choices, the test is aborted.

### **Supplemental Results**

Marker Gesture

| | Age | Rearing | Experience | Age + Experience | Age + Rearing | Age + Exp. + Rear. |
|---|---|---|---|---|---|---|
| Age (Weeks) | 0.114$^{***}$ (0.023) | | | 0.111$^{**}$ (0.044) | 0.114$^{***}$ (0.023) | 0.109$^{**}$ (0.044) |
| Group = Home | | 0.256 (0.242) | | | 0.228 (0.241) | 0.229 (0.241) |
| Experience | | | 0.201$^{***}$ (0.046) | 0.008 (0.091) | | 0.011 (0.091) |
| Constant | 0.718$^{**}$ (0.317) | 2.077$^{***}$ (0.206) | 1.893$^{***}$ (0.149) | 0.747 (0.466) | 0.565 (0.353) | 0.607 (0.485) |
| Observations | 503 | 503 | 503 | 503 | 503 | 503 |
| Akaike Inf. Crit. | 876.116 | 901.442 | 882.977 | 878.109 | 877.236 | 879.220 |

*Table S2: Marker Gesture Model Comparisons. For this and all subsequent model comparison tables, the values reported for each fixed effect variable are their beta coefficients, and the values in parentheses indicate their respective standard errors. ***p<.001, **p<.01, *p<0.5.*

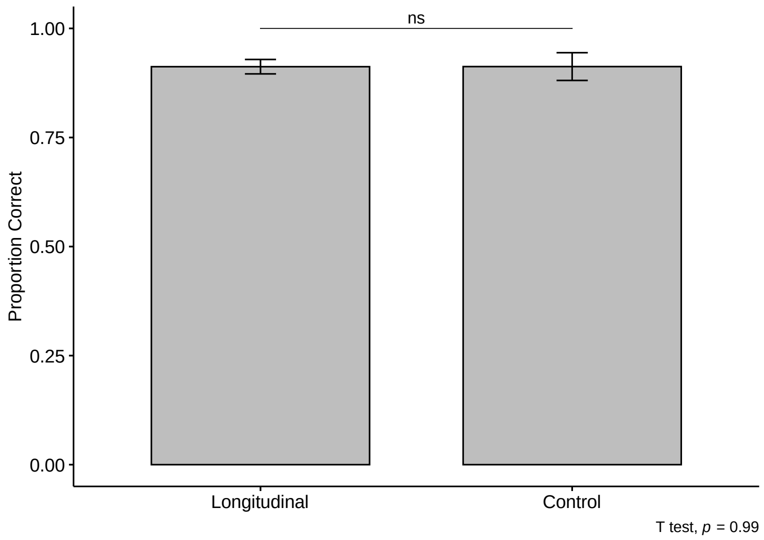

*Figure S1: Longitudinal vs Control Subjects' Performance on Marker Gesture Task at 17-21 weeks old*

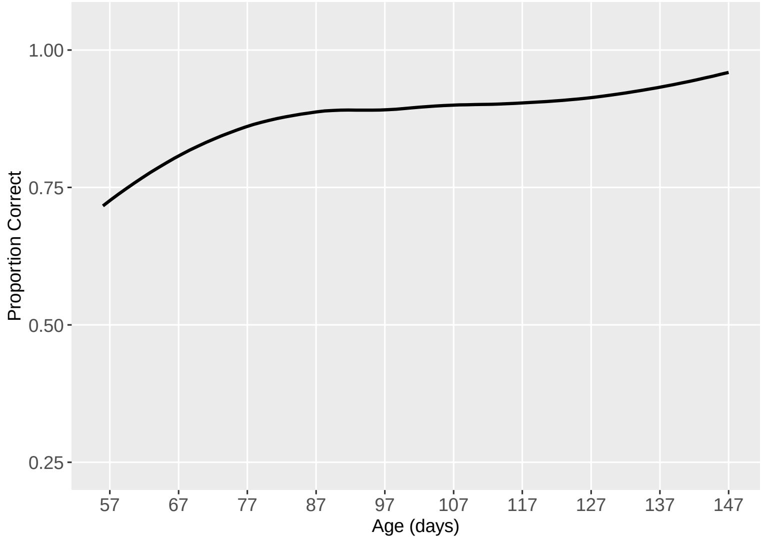

*Figure S2: Development of Marker Gesture Task by Age in Days*

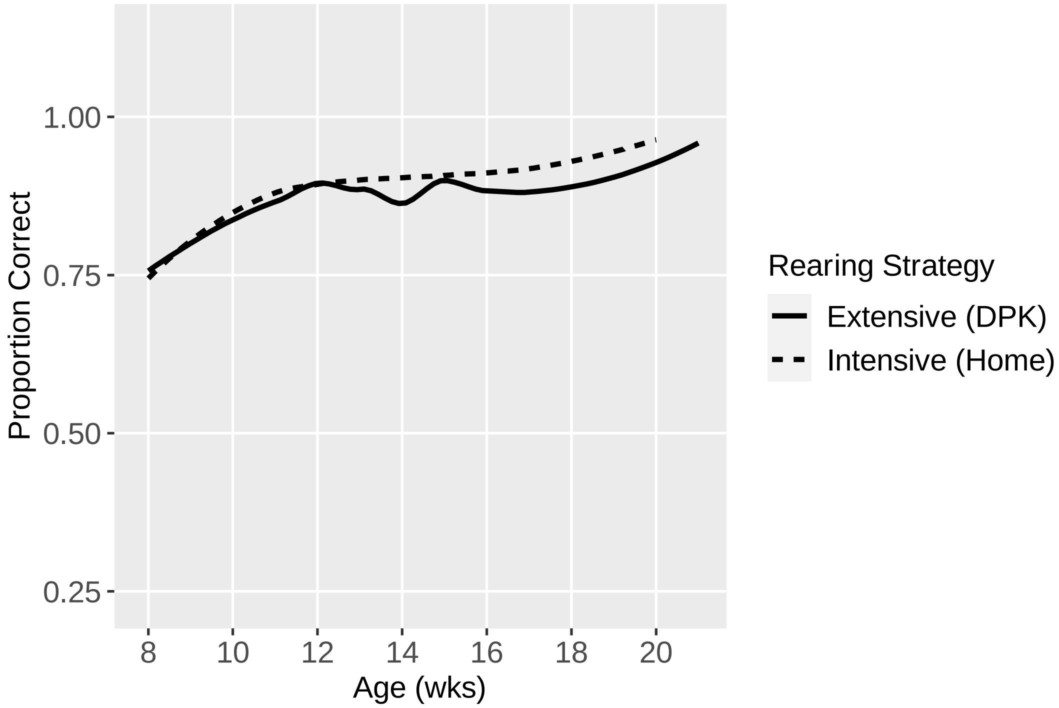

*Figure S3: Development of Marker Gesture Task by Rearing Strategy*

Working Memory (Delay)

| | Age | Rearing | Experience | Age + Experience | Age + Rearing | Age + Exp. + Rear. |
|---|---|---|---|---|---|---|
| Age (Weeks) | $0.067^{***}$ (0.001) | | | $0.201^{***}$ (0.051) | $0.067^{***}$ (0.001) | $0.204^{***}$ (0.052) |
| Group = Home | | $0.327^{***}$ (0.001) | | | $0.309^{***}$ (0.001) | 0.340 (0.253) |
| Experience | | | $0.067^{***}$ (0.001) | $-0.300^{***}$ (0.103) | | $-0.306^{***}$ (0.104) |
| Constant | $0.836^{***}$ (0.001) | $1.552^{***}$ (0.001) | $1.648^{***}$ (0.001) | -0.427 (0.523) | $0.632^{***}$ (0.001) | -0.679 (0.561) |
| Observations | 493 | 493 | 493 | 493 | 493 | 493 |
| Akaike Inf. Crit. | 1,080.790 | 1,091.217 | 1,089.886 | 1,072.844 | 1,081.248 | 1,073.020 |

*Table S3: Working Memory (Delay) Model Comparisons*

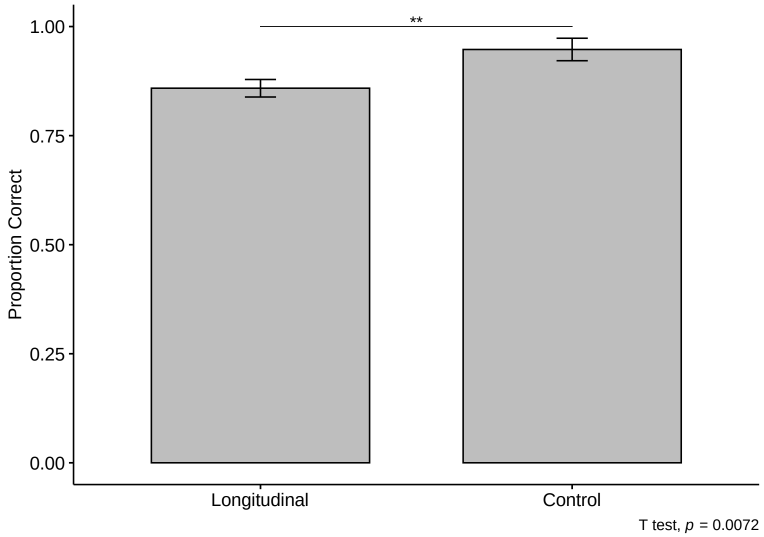

*Figure S4: Longitudinal vs Control Subjects' Performance on Working Memory (Delay) Task at 17-21 weeks old*

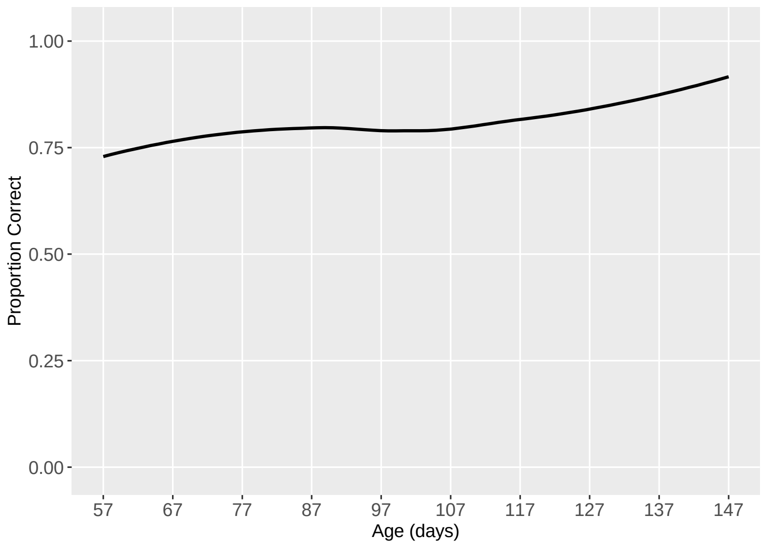

*Figure S5: Development of Working Memory (Delay) Task by Age in Days*

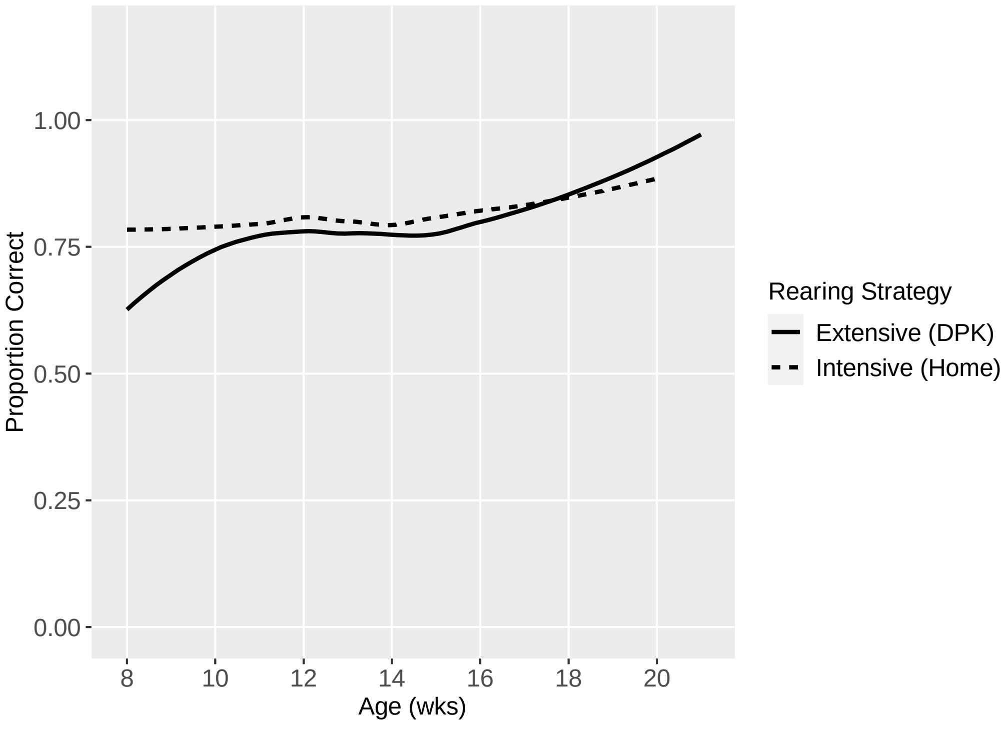

*Figure S6: Development of Working Memory (Delay) Task by Rearing Strategy*

#### Working Memory (Distraction)

| | Age | Rearing | Experience | Age + Experience | Age + Rearing | Age + Exp. + Rear. |
|---|---|---|---|---|---|---|
| Age (Weeks) | $0.091^{***}$ (0.017) | | | $0.117^{***}$ (0.002) | $0.091^{***}$ (0.016) | $0.118^{***}$ (0.034) |
| Group = Home | | -0.279 (0.205) | | | -0.311 (0.204) | -0.317 (0.204) |
| Experience | | | $0.147^{***}$ (0.034) | $-0.062^{***}$ (0.002) | | -0.061 (0.069) |
| Constant | -0.128 (0.243) | $1.306^{***}$ (0.172) | $0.846^{***}$ (0.117) | $-0.372^{***}$ (0.002) | 0.074 (0.277) | -0.164 (0.387) |
| Observations | 502 | 502 | 502 | 502 | 502 | 502 |
| Akaike Inf. Crit. | 1,223.128 | 1,251.835 | 1,234.305 | 1,224.399 | 1,222.845 | 1,224.050 |

*Table S4: Working Memory (Distraction) Model Comparisons*

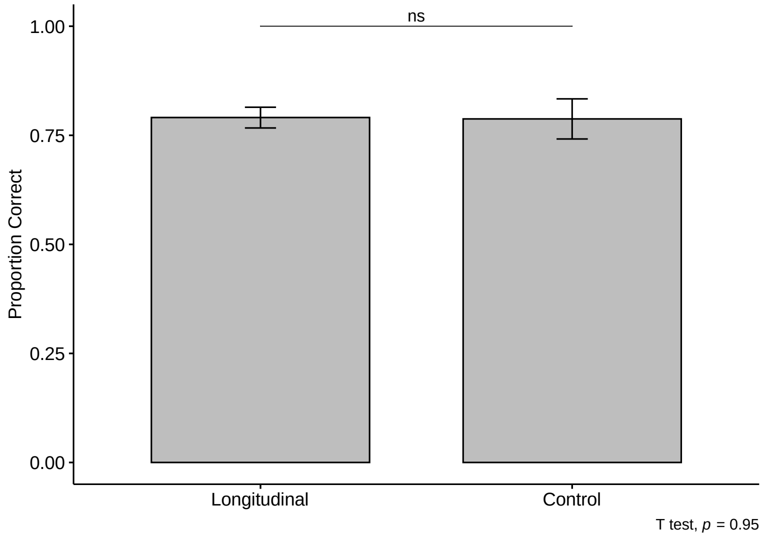

*Figure S7: Longitudinal vs Control Subjects' Performance on Working Memory (Distraction) Task at 17-21 weeks old*

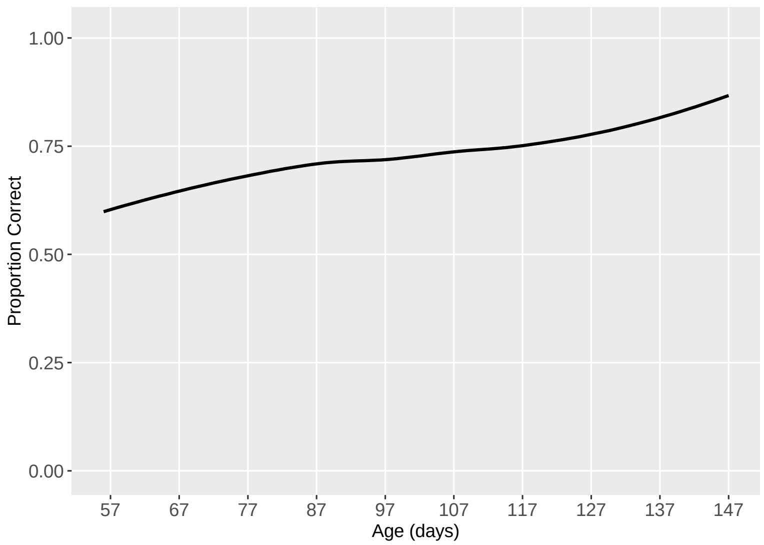

*Figure S8: Development of Working Memory (Distraction) Task by Age in Days*

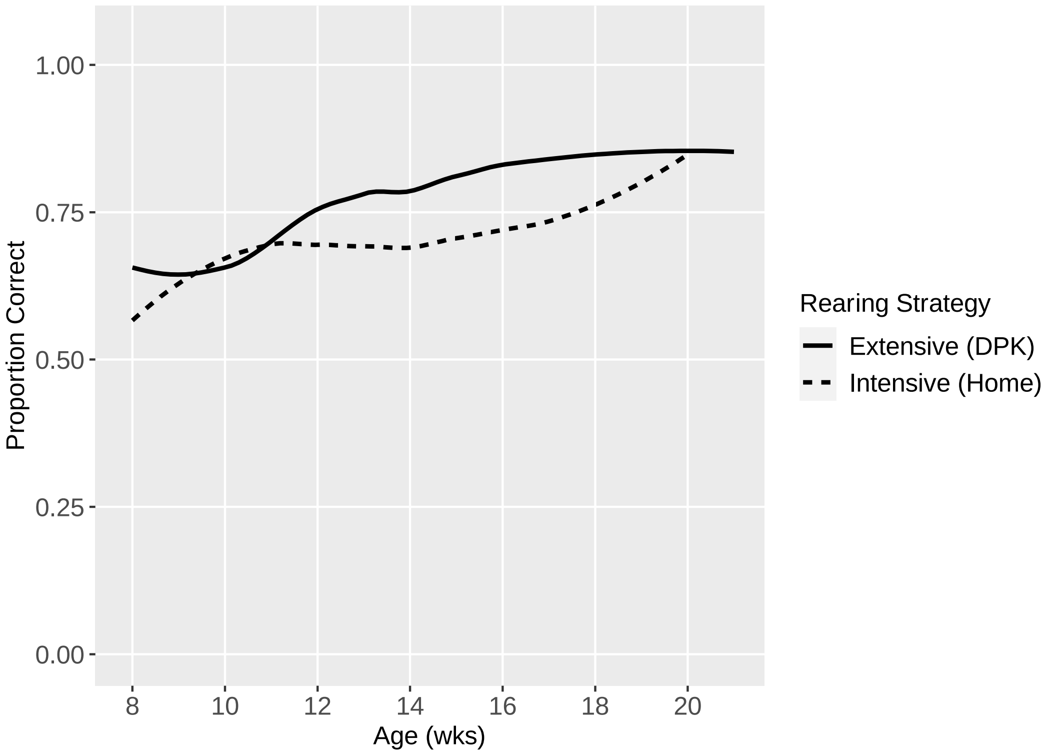

*Figure S9: Development of Working Memory (Distraction) by Rearing Strategy*

Auditory Discrimination

| | Age | Rearing | Experience | Age + Experience | Age + Rearing | Age + Exp. + Rear. |
|---|---|---|---|---|---|---|
| Age (Weeks) | -0.049*** (0.014) | | | 0.010 (0.024) | -0.049*** (0.014) | 0.009 (0.024) |
| Group = Home | | -0.159 (0.115) | | | -0.158 (0.116) | -0.151 (0.113) |
| Experience | | | -0.135*** (0.029) | -0.151*** (0.048) | | -0.150*** (0.048) |
| Constant | 1.328*** (0.209) | 0.750*** (0.097) | 0.930*** (0.083) | 0.828*** (0.260) | 1.434*** (0.224) | 0.934*** (0.272) |
| Observations | 497 | 497 | 497 | 497 | 497 | 497 |
| Akaike Inf. Crit. | 1,284.916 | 1,294.945 | 1,274.966 | 1,276.793 | 1,285.079 | 1,277.013 |

*Table S5: Auditory Discrimination Model Comparisons*

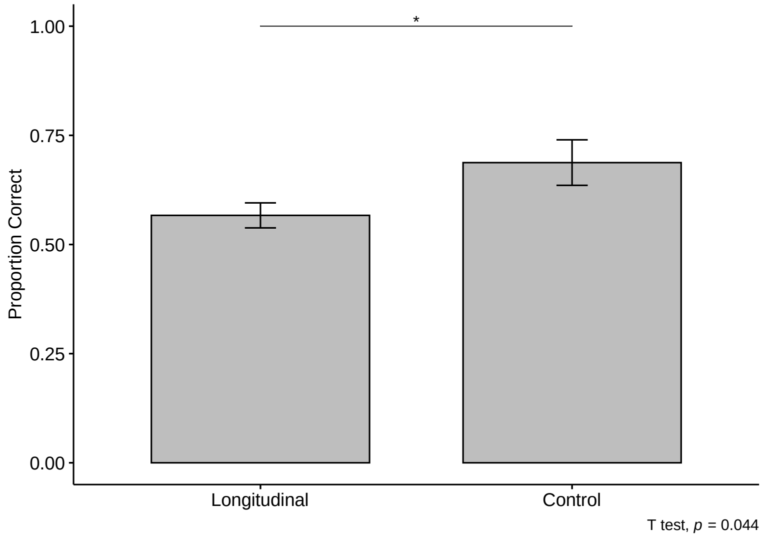

*Figure S10: Longitudinal vs Control Subjects' Performance on Auditory Discrimination Task at 17-21 weeks old*

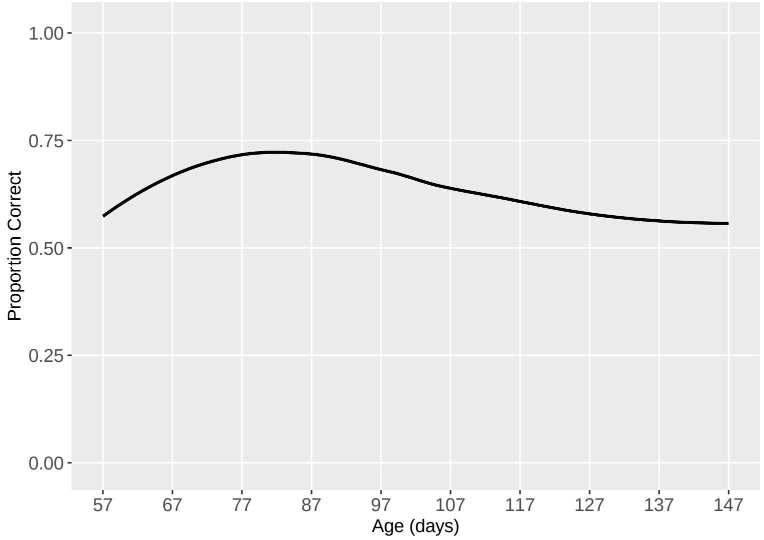

*Figure S11: Development of Auditory Discrimination Task by Age in Days*

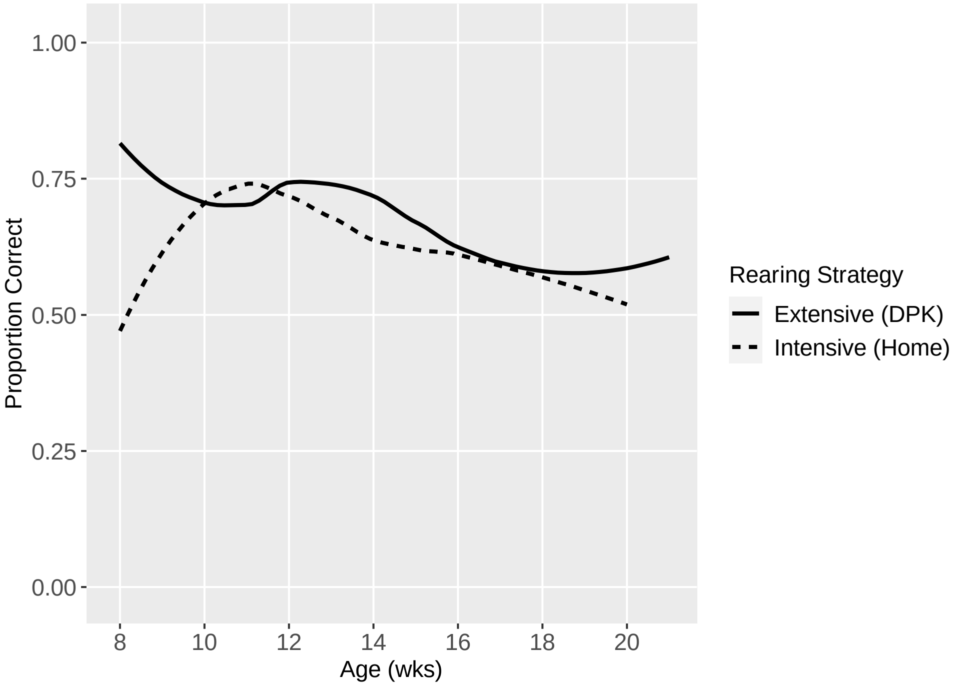

*Figure S12: Development of Auditory Discrimination Task by Rearing Strategy*

Odor Discrimination

| | Age | Rearing | Experience | Age + Experience | Age + Rearing | Age + Exp. + Rear. |
|---|---|---|---|---|---|---|
| Age (Weeks) | -0.029** (0.013) | | | -0.016 (0.021) | -0.029** (0.013) | -0.015 (0.021) |
| Group = Home | | 0.146 (0.099) | | | 0.146 (0.099) | 0.149 (0.099) |
| Experience | | | -0.062** (0.027) | -0.037 (0.043) | | -0.038 (0.043) |
| Constant | 0.513*** (0.191) | 0.003 (0.082) | 0.236*** (0.074) | 0.402* (0.231) | 0.412** (0.203) | 0.294 (0.242) |
| Observations | 492 | 492 | 492 | 492 | 492 | 492 |
| Akaike Inf. Crit. | 1,225.832 | 1,228.529 | 1,225.672 | 1,227.092 | 1,225.646 | 1,226.833 |

*Table S6: Odor Discrimination Task Model Comparisons*

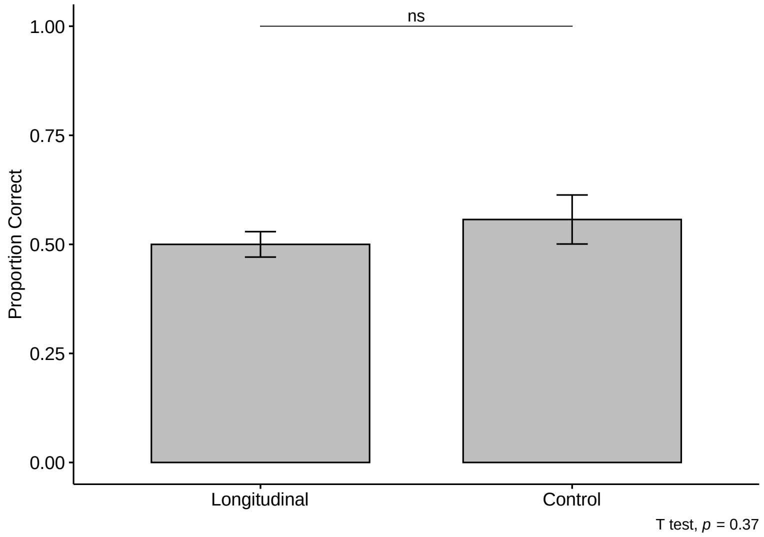

*Figure S13: Longitudinal vs Control Subjects' Performance on Odor Discrimination Task at 17-21 weeks old*

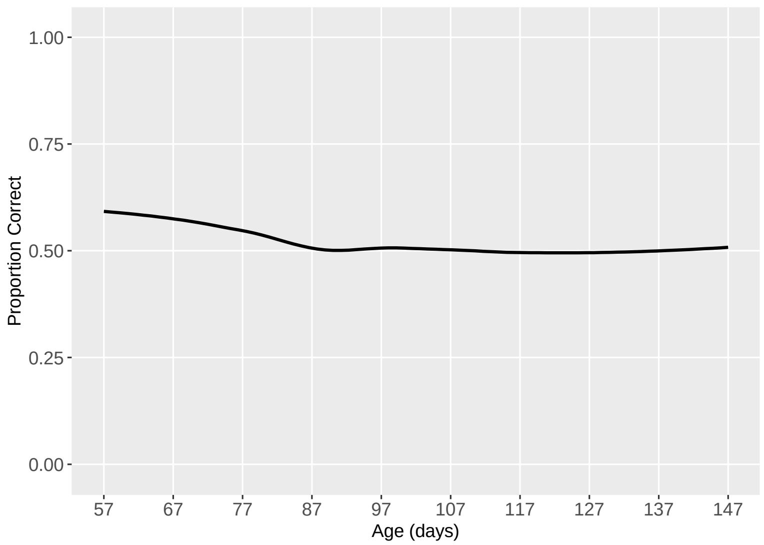

*Figure S14: Odor Discrimination Development by Age in Days*

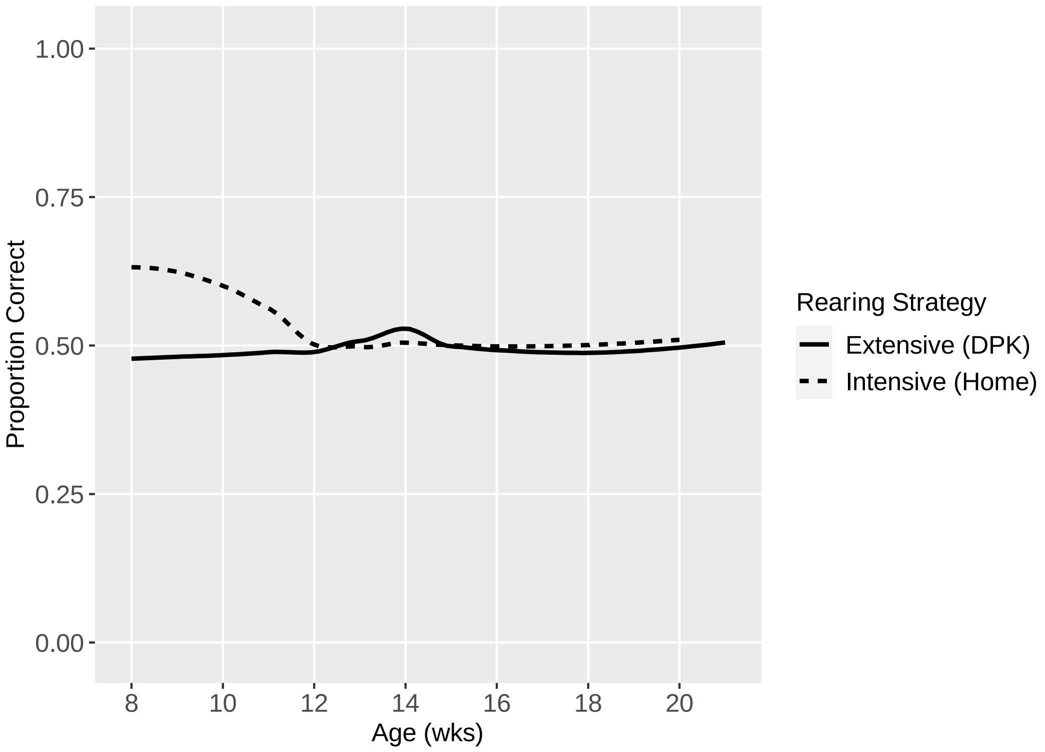

*Figure S15: Development of Odor Discrimination Task by Rearing Strategy*

Cylinder

| | Age | Rearing | Experience | Age + Experience | Age + Rearing | Age + Exp. + Rear. |
|---|---|---|---|---|---|---|
| Age (Weeks) | 0.425$^{***}$ (0.032) | | | 0.084$^{***}$ (0.028) | 0.426$^{***}$ (0.032) | 0.084$^{***}$ (0.028) |
| Group = Home | | 0.083 (0.143) | | | -0.065 (0.291) | 0.054 (0.175) |
| Experience | | | 0.524$^{***}$ (0.033) | 0.462$^{***}$ (0.038) | | 0.462$^{***}$ (0.038) |
| Constant | -3.944$^{***}$ (0.426) | 1.476$^{***}$ (0.118) | 0.148 (0.100) | -0.758$^{**}$ (0.310) | -3.906$^{***}$ (0.456) | -0.792$^{**}$ (0.330) |
| Observations | 498 | 498 | 498 | 498 | 498 | 498 |
| Akaike Inf. Crit. | 1,089.067 | 1,402.706 | 964.604 | 957.156 | 1,091.017 | 959.063 |

*Table S7: Cylinder Inhibitory Control Model Comparisons*

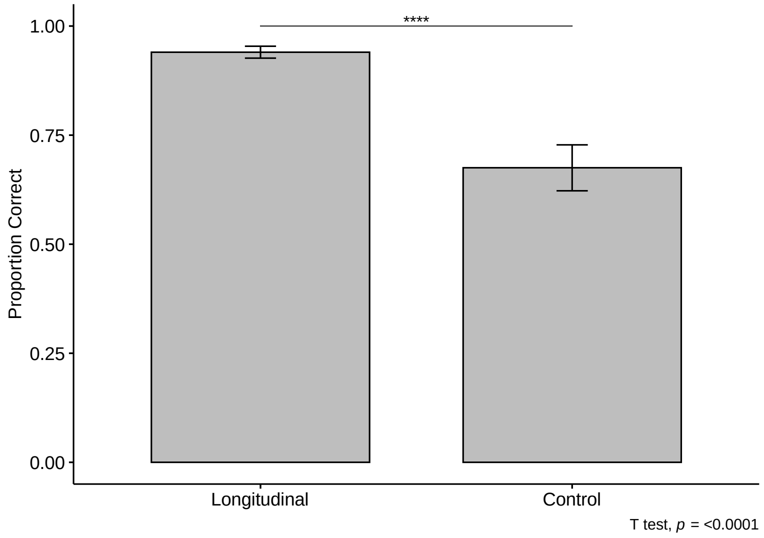

*Figure S16: Longitudinal vs Control Subjects' Performance on Cylinder Inhibitory Control Task at 17-21 weeks old*

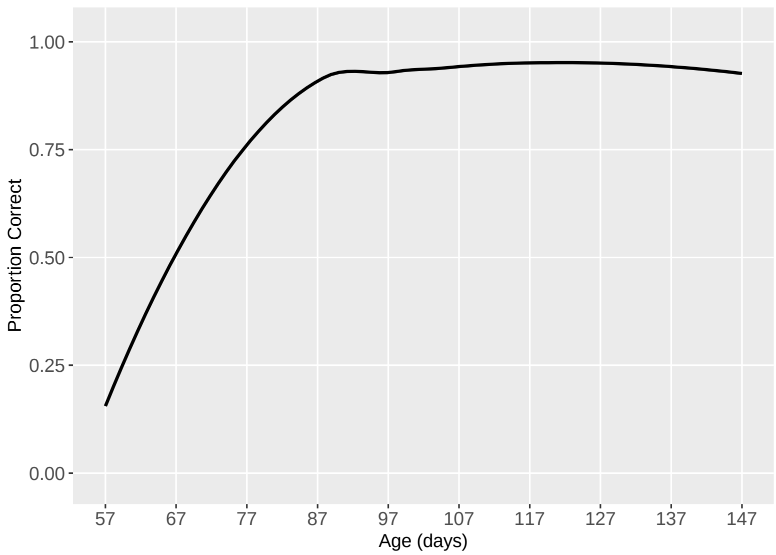

*Figure S17: Development of Cylinder Inhibitory Control Task by Age in Days*

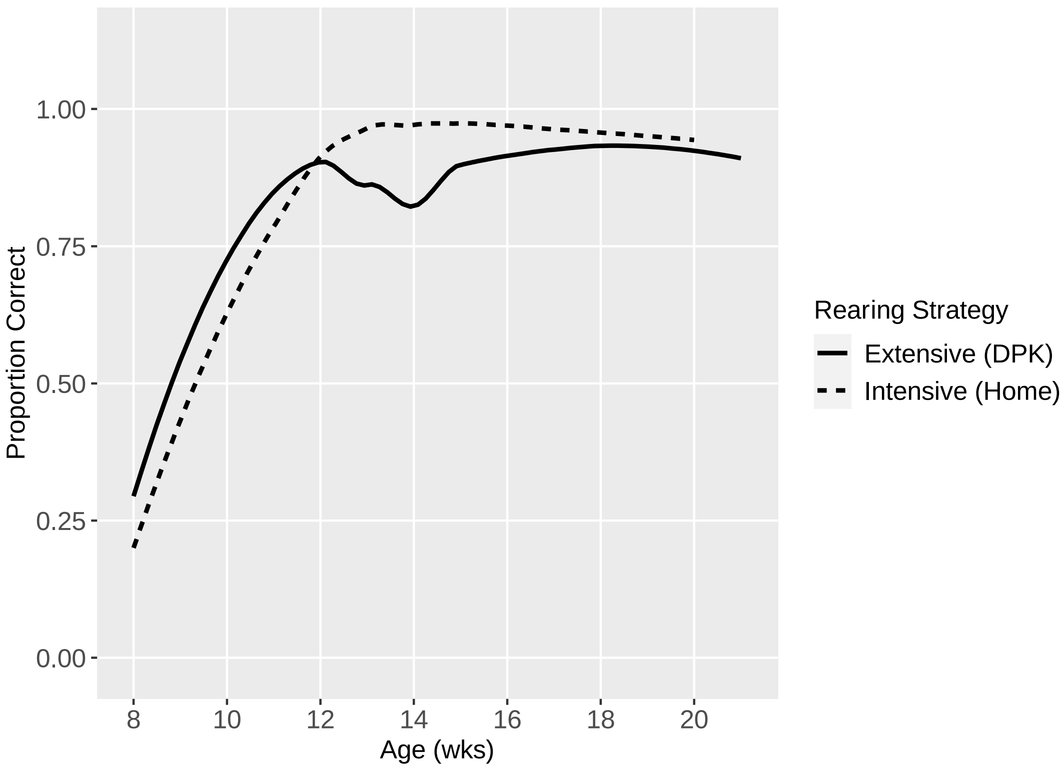

*Figure S18: Cylinder Inhibitory Control Task by Rearing Strategy*

| | Age | Rearing | Experience | Age + Experience | Age + Rearing | Age + Exp. + Rear. |
|---|---|---|---|---|---|---|
| Age (Weeks) | 0.231*** (0.017) | | | 0.057** (0.026) | 0.230*** (0.017) | 0.057** (0.026) |
| Group = Home | | 0.140 (0.124) | | | 0.109 (0.165) | 0.131 (0.139) |
| Experience | | | 0.256*** (0.017) | 0.209*** (0.026) | | 0.209*** (0.026) |
| Constant | -3.589*** (0.264) | -0.375*** (0.104) | -1.632*** (0.111) | -2.186*** (0.277) | -3.653*** (0.285) | -2.276*** (0.293) |
| Observations | 501 | 501 | 501 | 501 | 501 | 501 |
| Akaike Inf. Crit. | 1,426.717 | 1,649.291 | 1,366.060 | 1,363.344 | 1,428.290 | 1,364.459 |

*Table S8: Cylinder Reversal Learning Model Comparisons*

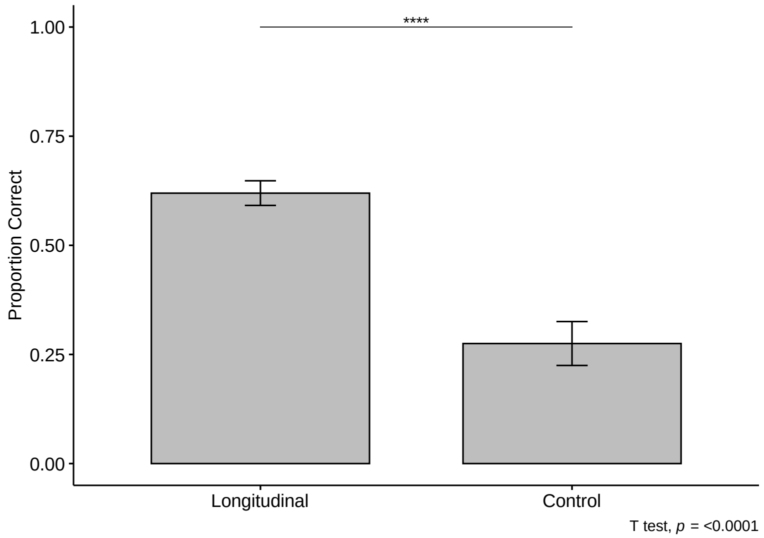

*Figure S19: Longitudinal vs Control Subjects' Performance on Cylinder Reversal Learning Task at 17-21 weeks old*

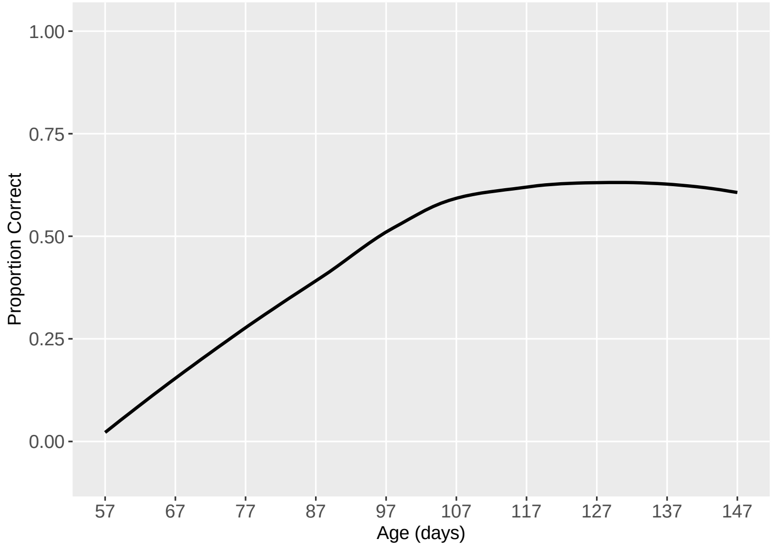

*Figure S20: Development of Cylinder Reversal Learning Task by Age in Days*

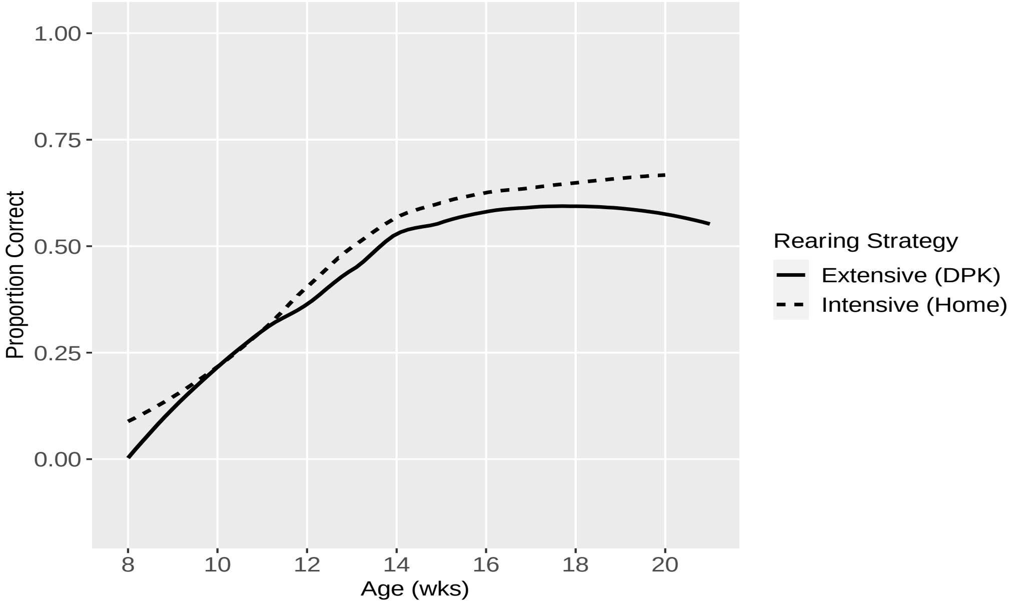

*Figure S21: Development of Cylinder Reversal Learning Task by Rearing Strategy*

Unsolvable (Eye Contact)

| | Age | Rearing | Experience | Age + Experience | Age + Rearing | Age + Exp. + Rear. |
|---|---|---|---|---|---|---|
| Age (Weeks) | 0.419$^{***}$ (0.043) | | | 0.194$^{***}$ (0.075) | 0.420$^{***}$ (0.043) | 0.195$^{***}$ (0.076) |
| Group = Home | | -0.091 (0.452) | | | -0.216 (0.457) | -0.196 (0.449) |
| Experience | | | 0.876$^{***}$ (0.086) | 0.550$^{***}$ (0.152) | | 0.550$^{***}$ (0.153) |
| Constant | -2.272$^{***}$ (0.631) | 3.601$^{***}$ (0.374) | 1.746$^{***}$ (0.275) | -0.281 (0.831) | -2.138$^{***}$ (0.698) | -0.154 (0.881) |
| Observations | 498 | 498 | 498 | 498 | 498 | 498 |
| Akaike Inf. Crit. | 2,645.914 | 2,726.166 | 2,638.162 | 2,636.923 | 2,647.425 | 2,638.499 |

*Table S9: Unsolvable (Total Looking Time) Model Comparisons*

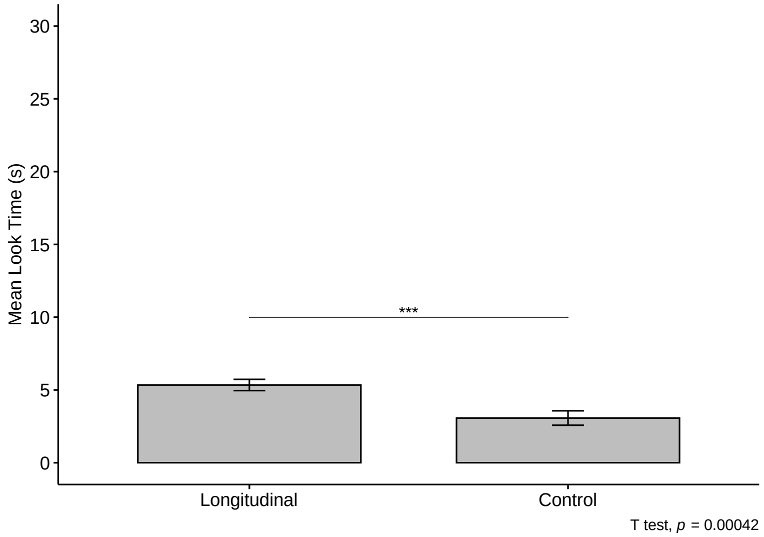

*Figure S22: Longitudinal vs Control Subjects' Performance on Unsolvable Task (Looking Time) at 17-21 weeks old*

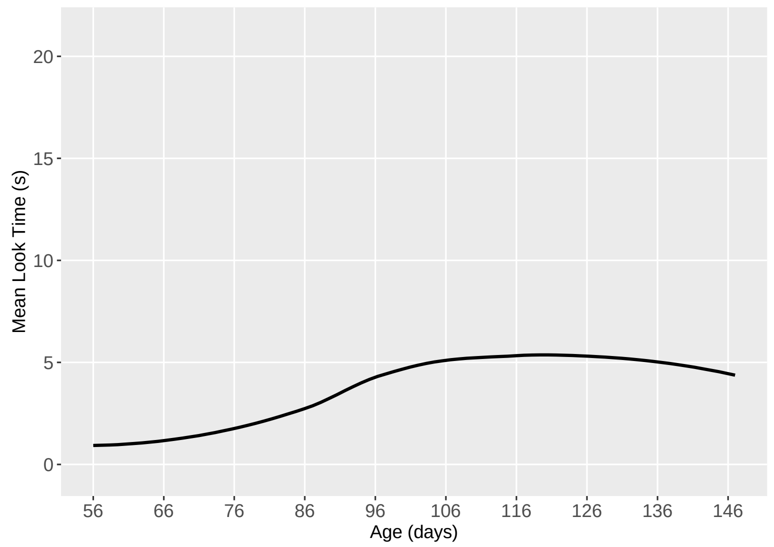

*Figure S23: Development of Unsolvable Task (Looking Time) by Age in Days*

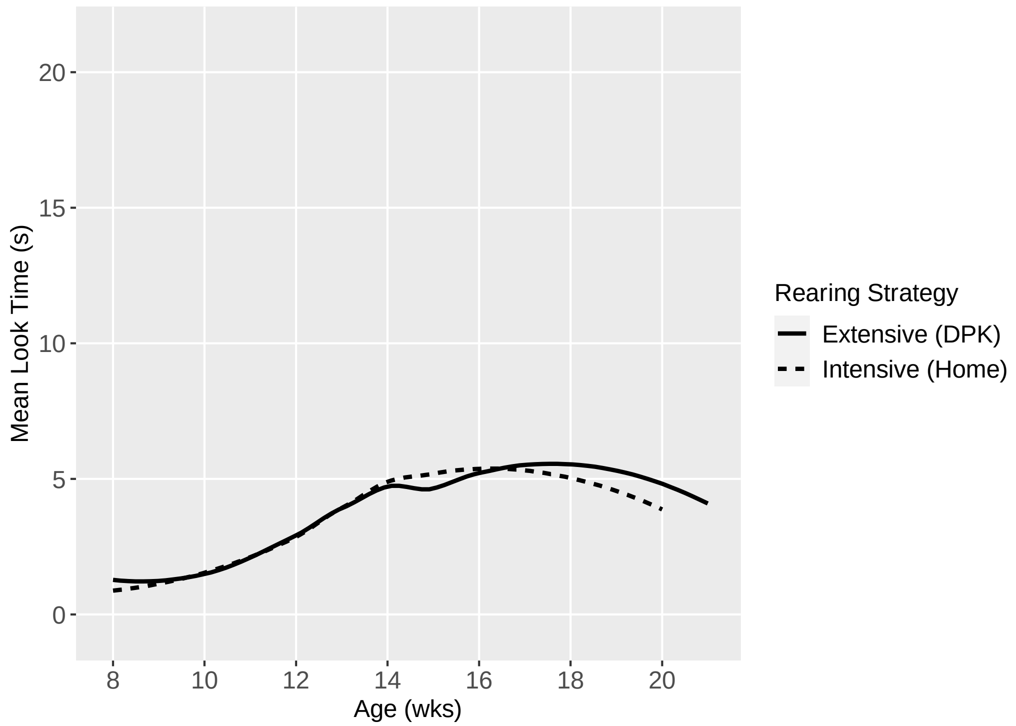

*Figure S24: Development of Unsolvable Task (Looking Time) by Rearing Strategy*

| | Age | Rearing | Experience | Age + Experience | Age + Rearing | Age + Exp. + Rear. |
|---|---|---|---|---|---|---|
| Age (Weeks) | 0.179*** (0.017) | | | 0.081*** (0.031) | 0.179*** (0.017) | 0.080*** (0.031) |
| Group = Home | | 0.182 (0.191) | | | 0.117 (0.210) | 0.142 (0.203) |
| Experience | | | 0.374*** (0.035) | 0.234*** (0.063) | | 0.235*** (0.063) |
| Constant | -1.691*** (0.250) | 0.622*** (0.157) | 0.071 (0.113) | -0.777** (0.341) | -1.766*** (0.285) | -0.864** (0.362) |
| Observations | 498 | 498 | 498 | 498 | 498 | 498 |
| Akaike Inf. Crit. | 1,442.517 | 1,561.652 | 1,435.963 | 1,431.142 | 1,444.211 | 1,432.655 |

*Table S10: Unsolvable Task (Proportion of Trials Looked) Model Comparisons*

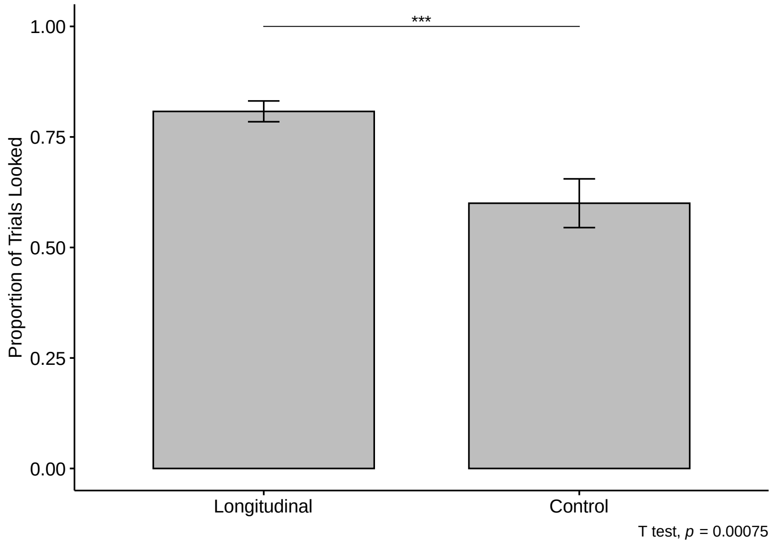

*Figure S25: Longitudinal vs Control Subjects' Performance on Unsolvable Task (Proportion of Trials Looked) at 17-21 weeks old*

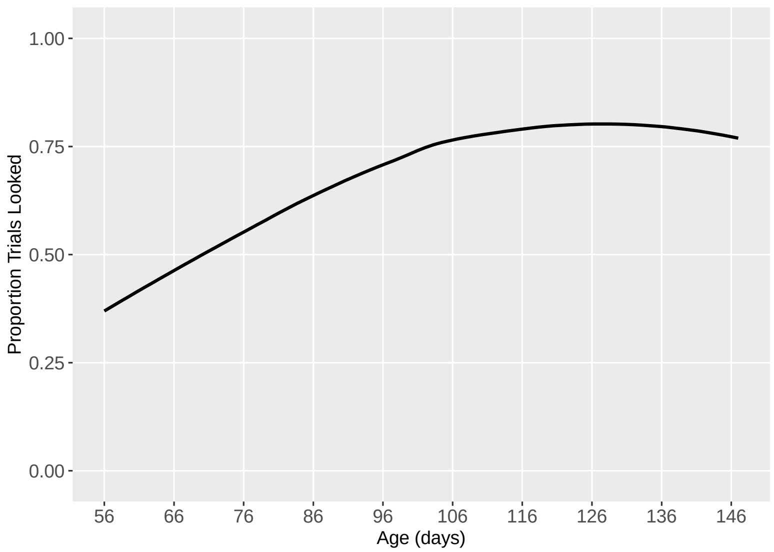

*Figure S26: Development of Unsolvable Task (Proportion of Trials Looked) by Age in Days*

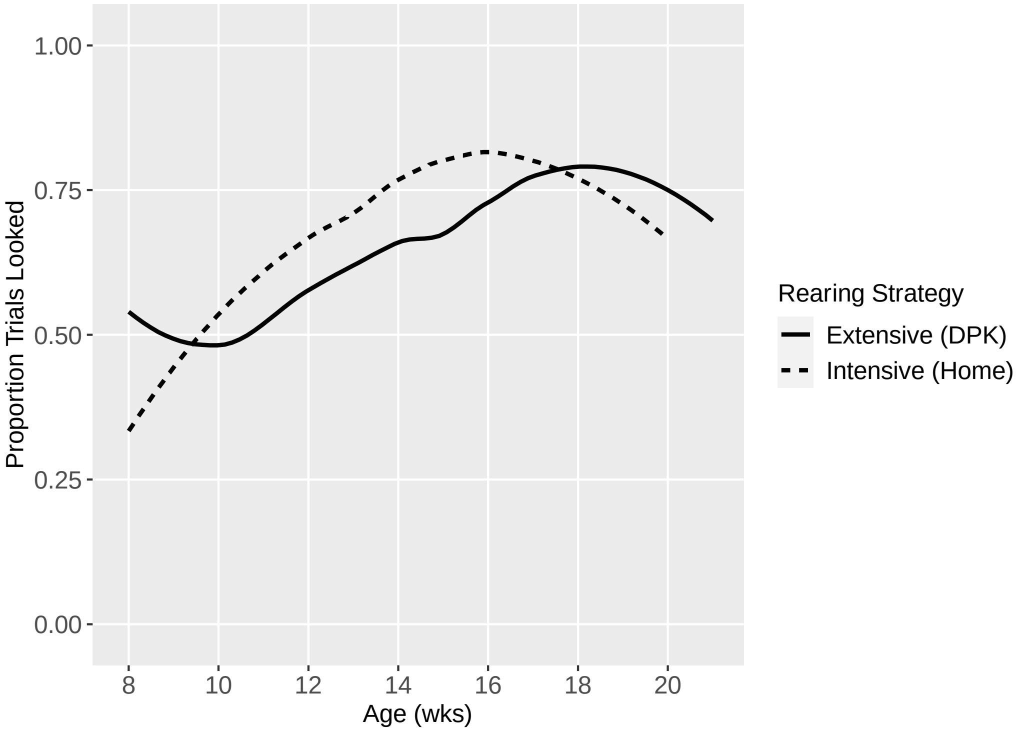

*Figure S27: Development of Unsolvable Task (Proportion of Trials Looked) by Rearing Strategy*

Causal Reasoning

| | Age | Rearing | Experience | Age + Experience | Age + Rearing | Age + Exp. + Rear. |
|---|---|---|---|---|---|---|
| Age (Weeks) | $0.191^{***}$ (0.018) | | | 0.006 (0.025) | $0.191^{***}$ (0.018) | 0.004 (0.024) |
| Group = Home | | $-0.256^{*}$ (0.132) | | | $-0.339^{*}$ (0.174) | $-0.354^{**}$ (0.143) |
| Experience | | | $0.476^{***}$ (0.035) | $0.467^{***}$ (0.053) | | $0.472^{***}$ (0.052) |
| Constant | $-2.071^{***}$ (0.260) | $0.741^{***}$ (0.112) | $-0.366^{***}$ (0.090) | -0.429 (0.272) | $-1.825^{***}$ (0.279) | -0.161 (0.290) |
| Observations | 473 | 473 | 473 | 473 | 473 | 473 |
| Akaike Inf. Crit. | 1,246.048 | 1,378.505 | 1,169.267 | 1,171.207 | 1,244.197 | 1,167.057 |

*Table S11: Causal Reasoning Model Comparisons*

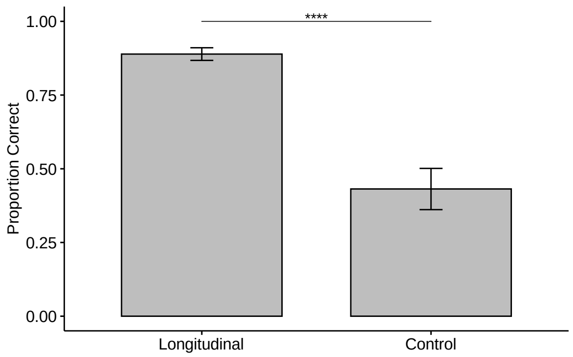

*Figure S28: Longitudinal vs Control Subjects' Performance on Causal Reasoning Task at 17-21 weeks old*

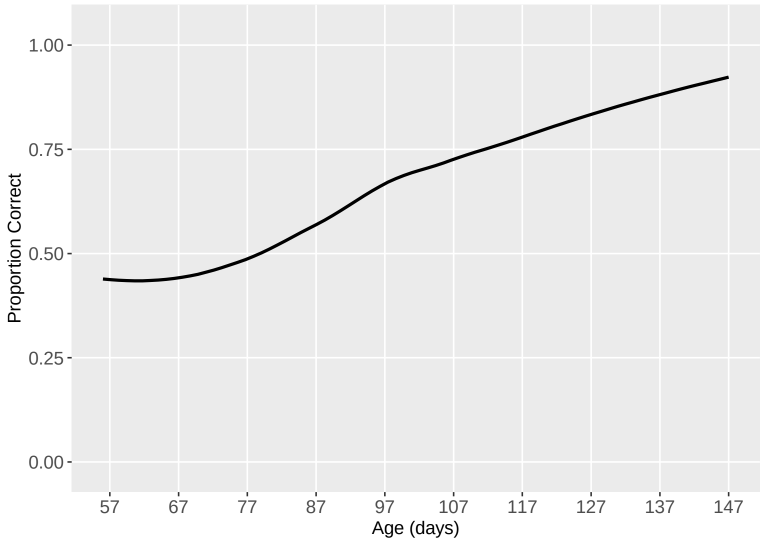

*Figure S29: Development of Causality Task by Age in Days*

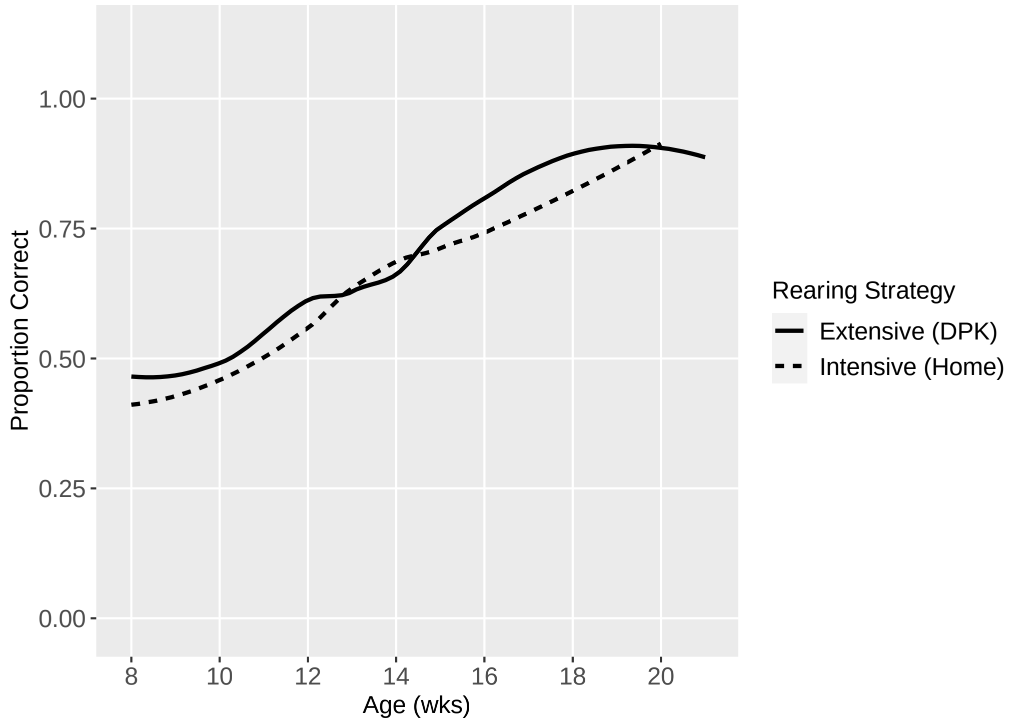

*Figure S30: Development of Causality by Rearing Strategy*

<u>Momentary Pointing</u>

| | Age | Rearing | Experience | Age + Experience | Age + Rearing | Age + Exp. + Rear. |
|---|---|---|---|---|---|---|
| Age (Weeks) | -0.007 (0.013) | | | -0.023 (0.021) | -0.007 (0.013) | -0.023 (0.021) |
| Group = Home | | 0.075 (0.098) | | | 0.075 (0.098) | 0.073 (0.098) |
| Experience | | | 0.006 (0.027) | 0.043 (0.043) | | 0.042 (0.043) |
| Constant | 0.109 (0.189) | -0.043 (0.081) | -0.003 (0.073) | 0.240 (0.231) | 0.057 (0.201) | 0.187 (0.241) |
| Observations | 494 | 494 | 494 | 494 | 494 | 494 |
| Akaike Inf. Crit. | 1,268.720 | 1,268.422 | 1,268.967 | 1,269.728 | 1,270.127 | 1,271.171 |

*Table S12: Momentary Pointing Model Comparisons*

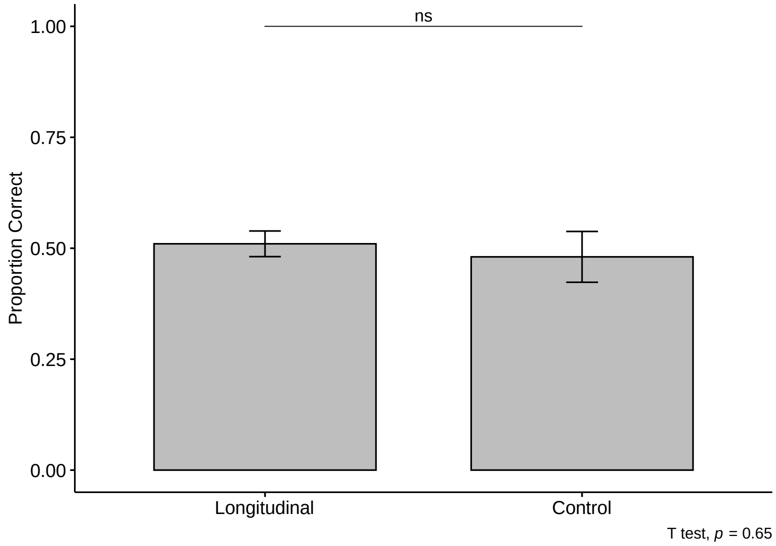

*Figure S31: Longitudinal vs Control Subjects' Performance on Momentary Pointing Task at 17-21 weeks old*

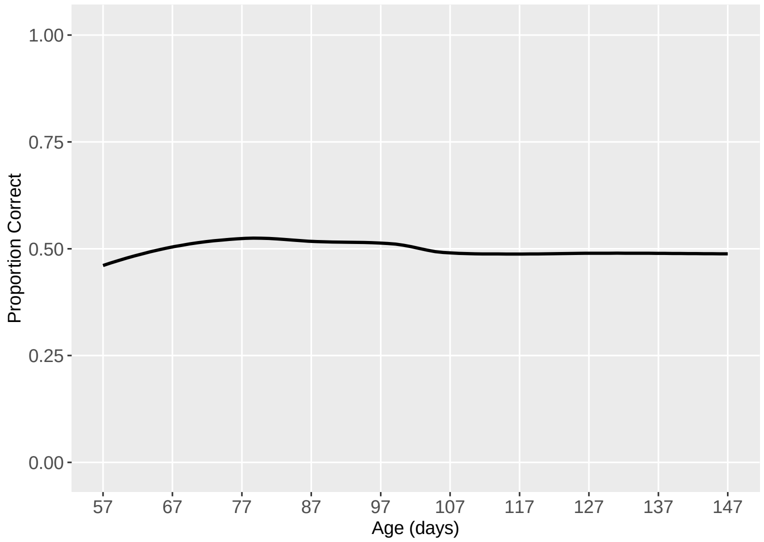

*Figure S32: Development of Momentary Pointing Task by Age in Days*

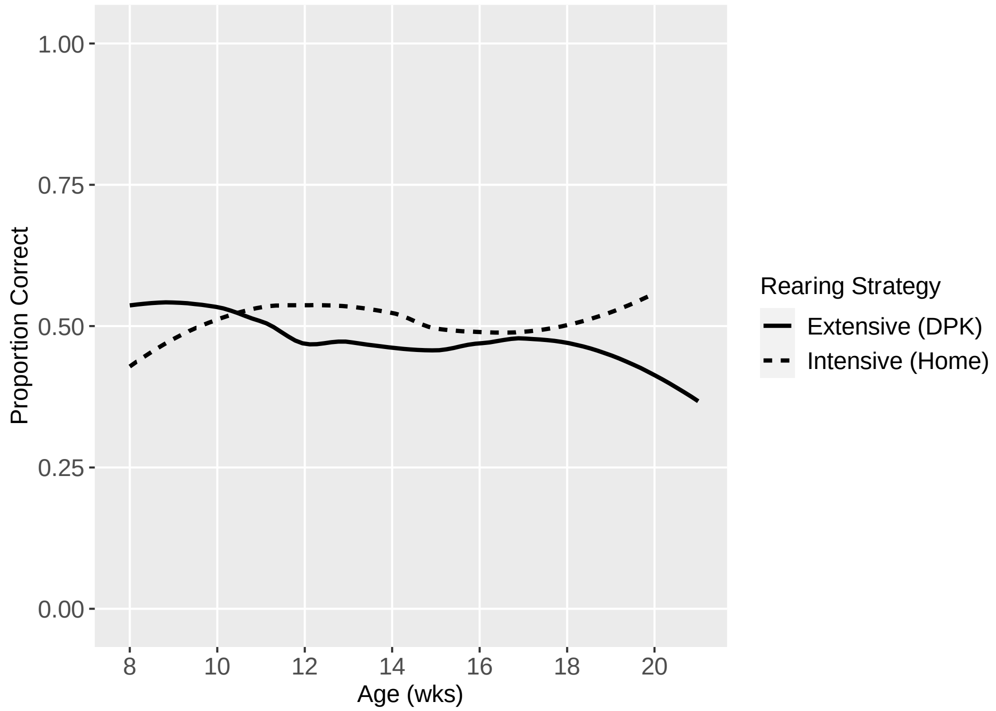

*Figure S33: Development of Momentary Pointing Task by Rearing Strategy*

Breakpoint Regressions By Task

*Marker Gesture*

Segmented regressions fit to the data yielded two segments (slope 1 = 0.0073, SE = 0.0029; slope 2 = 0.00079, SE = 0.00053) with a breakpoint identified at 76.85 days of age (Figure ).

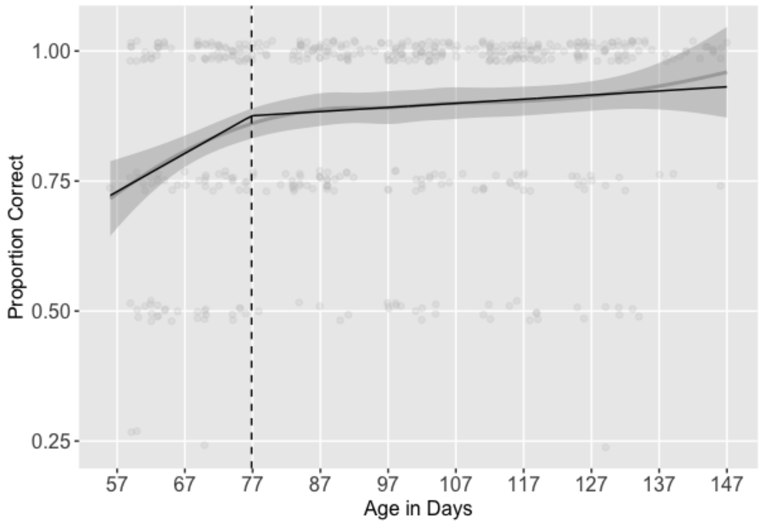

*Figure S34: Segmented regression lines (black) fit to Marker Gesture task development curve (gray); breakpoint indicated by dashed line*

*Working Memory (Delay)*

Segmented regressions fit to the data yielded two segments (slope 1 = 0.0011, SE = 0.00051; slope 2 = 0.0091, SE = 0.0093) with a breakpoint identified at 134.00 days of age (Figure ).

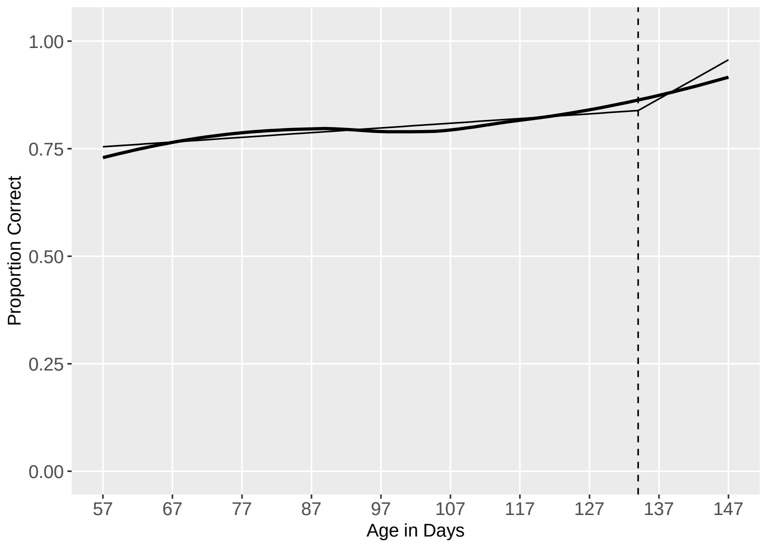

*Figure S35: Segmented regression lines (black) fit to Working Memory (Delay) task development curve (gray); breakpoint indicated by dashed line*

<u>*Working Memory (Distraction)*</u>

Segmented regressions fit to the data yielded two segments (slope 1 = 0.0020, SE = 0.00055; slope 2 = 0.0064, SE = 0.0061) with a breakpoint identified at 129.00 days of age (Figure ).

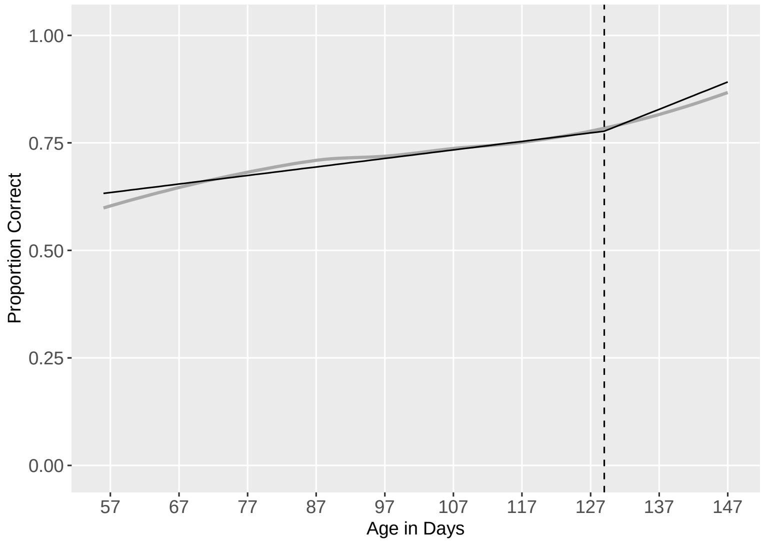

*Figure S36: Segmented regression lines (black) fit to Working Memory (Distraction) task development curve (gray); breakpoint indicated by dashed line*

*Cylinder Inhibitory Control*

Segmented regressions fit to the data yielded two segments (slope 1 = 0.028, SE = 0.0028; slope 2 = 0.00072, SE = 0.00067) with a breakpoint identified at 82.38 days of age (Figure S37).

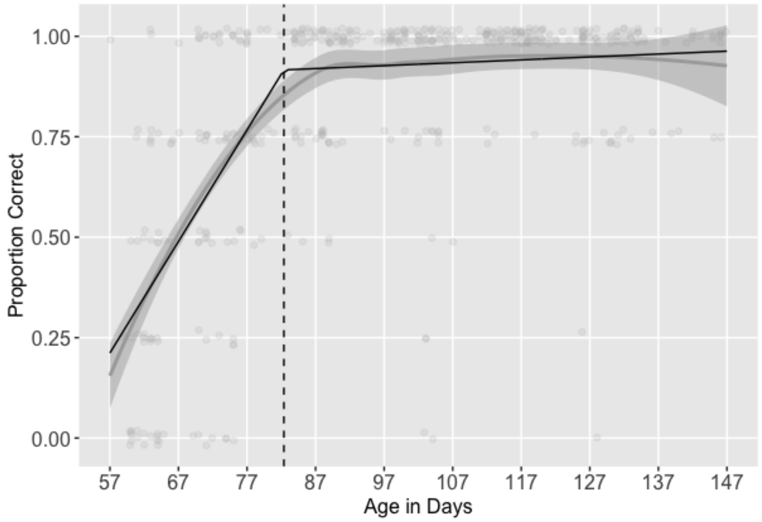

*Figure S37: Segmented regression lines (black) fit to Cylinder Inhibitory Control task development curve (gray); breakpoint indicated by dashed line*

*Cylinder Reversal Learning*

Segmented regressions fit to the data yielded two segments (slope 1 = 0.012, SE = 0.0013; slope 2 = 0.0017, SE = 0.0014) with a breakpoint identified at 101.06 days of age (Figure S38).

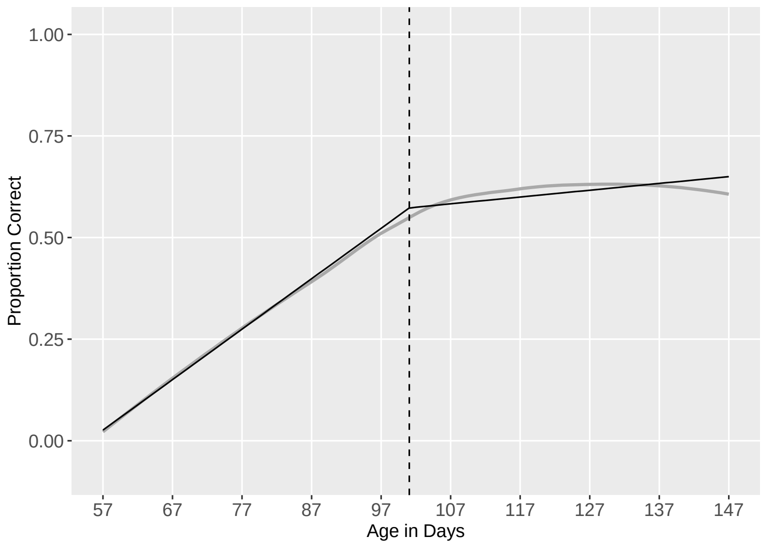

*Figure S38: Segmented regression lines (black) fit to Cylinder Reversal Learning task development curve (gray); breakpoint indicated by dashed line*

<u>*Eye Contact (Unsolvable) Proportion Trials Looked*</u>

Segmented regressions fit to the data yielded two segments (slope 1 = 0.0079, SE = 0.0012; slope 2 = -0.00019, SE = 0.0023) with a breakpoint identified at 107.64 days of age (Figure S39).

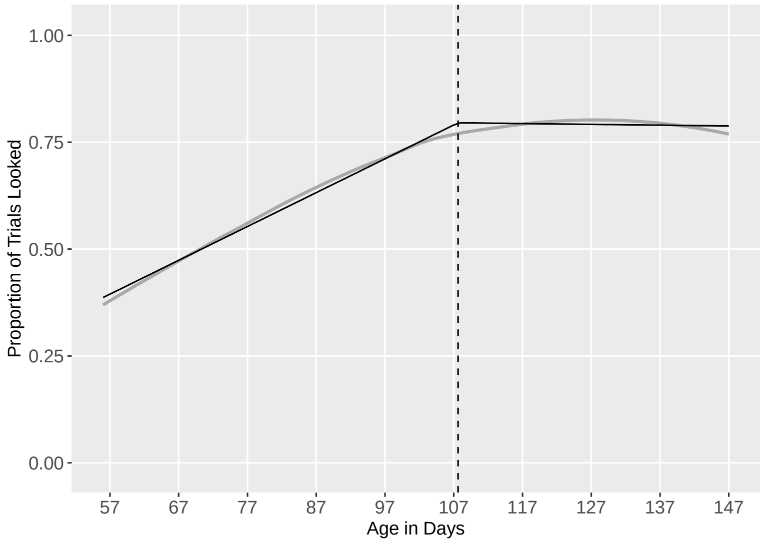

*Figure S39: Segmented regression lines (black) fit to Unsolvable (Proportion of Trials Looked) task development curve (gray); breakpoint indicated by dashed line*

*Eye Contact (Unsolvable) Mean Looking Time*

Segmented regressions fit to the data yielded two segments (slope 1 = 0.10, SE = 0.014; slope 2 = -0.0020, SE = 0.026) with a breakpoint identified at 106.00 days of age (Figure S40).

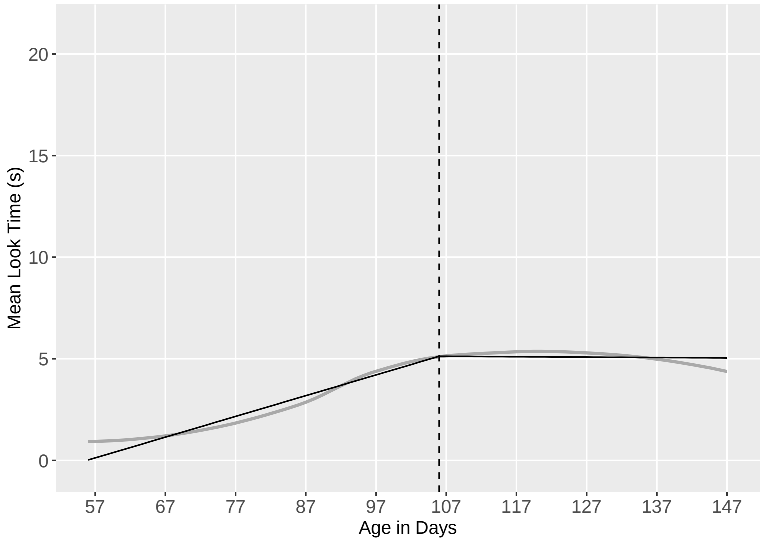

*Figure S40: Segmented regression lines (black) fit to Unsolvable (Mean Looking Time) task development curve (gray); breakpoint indicated by dashed line*

*Causal Reasoning*

Segmented regressions fit to the data yielded two segments (slope 1 = -0.055, SE = 0.031; slope 2 = 0.0069, SE = 0.00052) with a breakpoint identified at 62.00 days of age (Figure ).

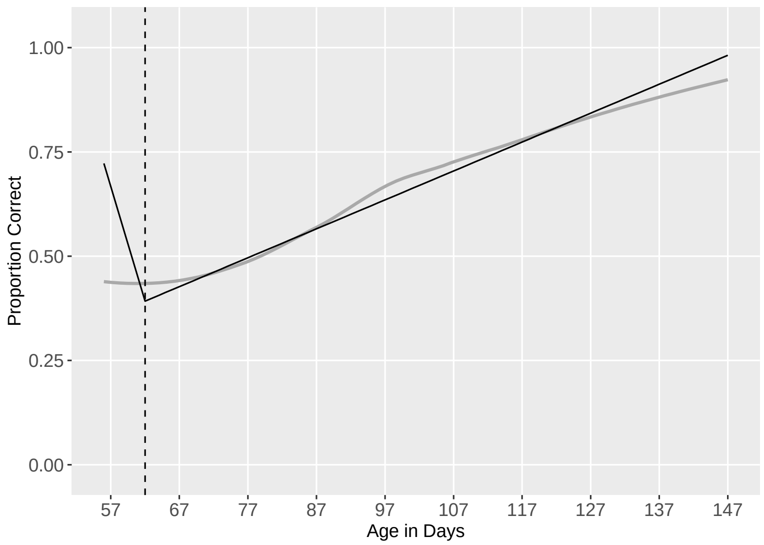

*Figure S41: Segmented regression lines (black) fit to Causal Reasoning task development curve (gray); breakpoint indicated by dashed line*

| Task | Longitudinal Last Test Mean (only those >=17 wks) | Control Mean (only those >=17 wks) | Adult Mean | Source of Adult Mean |
|---|---|---|---|---|
| Marker Gesture % correct | 91% | 91% | 89.32% | Bray et al. 2020 (12 trials) |
| Working Memory (20 sec. Delay) % correct | 86% | 95% | 73.09% | Bray et al. 2021 (mean of 6 trials with each increasing delay, and 4 hiding places instead of 2) |
| Working Memory (Distraction) | 79% | 79% | NA | Maclean & Hare 2018 |
| Auditory Disc % correct | 57% | 69% | 65.47 | Bray et al. 2021 (8 trials) |
| Odor Disc % correct | 50% | 56% | 60.94 | Bray et al. 2021 (8 trials) |
| Inhibitory Control (Cylinder Detour)* % correct | 94% | 68% | 75.94% | Bray et al 2021 (8 trials) |
| Reversal Learning (Cylinder | 62% | 28% | 59.59% | Bray et al 2021 (8 trials) |

| | | | | |
|---|---|---|---|---|
| Reversal) % correct | | | | |
| Unsolvable Eye Contact (looking time in seconds out of 30 second trial) | 5.34 sec | 3.07 sec | 3.3 sec | Bray et al. 2021 |
| Unsolvable Eye Contact (percent of trials made any eye contact)* | 81% | 60% | NA | Bray et al 2021 (not calculated) |
| Causal Reasoning | 89% | 43% | NA | Maclean & Hare 2018 |
| Momentary Pointing (% correct) | 51% | 50% | 70% | Salomons et al. 2024 (6 trials; pet dogs) |

*Table S13: Comparing Longitudinal Puppy, Control Puppy, and Adult Means*

| | 2 sessions | 3 sessions | 4 sessions | 5 sessions | 6 sessions |
|---|---|---|---|---|---|
| **8 weeks** | 0 | 4 | 1 | 7 | 20 |
| **9 weeks** | 0 | 0 | 0 | 7 | 23 |
| **10 weeks** | 0 | 0 | 3 | 7 | 10 |
| **11 weeks** | 0 | 0 | 0 | 0 | 1 |
| **12 weeks** | 3 | 0 | 0 | 0 | 0 |
| **13 weeks** | 0 | 0 | 0 | 0 | 0 |
| **14 weeks** | 0 | 1 | 4 | 0 | 0 |

*Table S14: Matrix of Total Number of Testing Sessions Completed (columns) by Age at First Testing Session (rows)*

| ***Session 1*** | ***Emergence Position*** |
|---|---|
| Unsolvable (Eye Contact) | 2 |
| Marker Gesture | 1 |
| Working Memory (Distraction) | 1 |
| Causal Reasoning | 3 |
| ***Session 2*** | |
| Cylinder (Inhibitory Control & Reversal Learning) | 2 & 4 |
| Momentary Pointing Gesture | 5 |
| Working Memory (Delay) | 1 |
| Auditory Discrimination | 1 |
| Odor Discrimination | 1 |

*Table S15: Order of test administration within the sessions compared to order of emergence across development. Emergence Positions: 1 = 8-9 wks, 2= 10-11 wks, 3 = 12-13 wks, 4 = 14-15 wks, 5 = does not emerge during the tested window but does emerge by adulthood*
